# Integrated Multi-omics Analyses of NFKB1 patients B cells points towards an up regulation of NF-κB network inhibitors

**DOI:** 10.1101/2022.11.22.517350

**Authors:** Nadezhda Camacho-Ordonez, Neftali Ramirez, Sara Posadas-Cantera, Andrés Caballero-Oyteza, Manfred Fliegauf, Fangwen Zhao, Maria Guarini, Victoria Gernedl, Mateo Pecoroaro, Klaus Warnatz, Christoph Bock, Esteban Ballestar, Roger Geiger, Michele Proietti, Bodo Grimbacher

## Abstract

The transcription factor NF-κB plays a pivotal role in the adaptive immune response. Pathogenic variants in *NFKB1* are the most common genetic etiology of common variable immunodeficiency (CVID). Patients frequently present with impaired terminal B cell differentiation, autoimmunity, and hyperinflammatory immune dysregulation. NF-κB signaling and target gene expression are expected to be dysregulated in *NFKB1*-mutated patients. Here, we performed a multi-omics characterization of B cells from a cohort of clinically affected and unaffected *NFKB1* mutation carriers. Our analysis identified specific epigenetic dysregulation and gene expression differences on B cells from *NFKB1*-mutated patients. We observed an aberrant expression of negative regulators of NF-κB signaling in *NFKB1* mutation carriers, which may be a key factor for the autoinflammatory phenotype of these patients. Moreover, our analysis points towards a dysregulation of *XBP1* and *BCL3*, key players of B cell activation and proliferation at different stages of B cell differentiation. The reduced expression of negative regulators of the NF-κB network is likely to be one of several mechanisms responsible for the aberrant NF-κB signaling, which impairs the maintenance of a normal humoral immune response. In summary, our findings highlight epigenetic and gene expression changes in B cells associated with *NFKB1* mutations. Our data give insight into future therapeutic opportunities for patients with *NFKB1* (haplo)insufficiency.

## Introduction

Heterozygous damaging variants in *NFKB1* constitute the most common genetic cause of common variable immunodeficiency (CVID)^1–8^. The NF-κB signaling pathway plays a central role in multiple biological processes, including the adaptive immune response^9,10^. NF-κB comprises five family members: p65 (RelA), RelB, c-Rel, p105/p50 (*NFKB1*, activated by the canonical pathway), and p100/p52 (*NFKB2*, activated by the non-canonical pathway)^11^. *NFKB1* encodes the precursor p105, which undergoes proteasomal processing of its C-terminal half to generate the mature p50. In the cytoplasm, p50 predominantly assembles with p65 (RelA) but remains inactive while forming a complex with the inhibitor of NF-κB-alpha (IκBα). Upon pathway stimulation, IκBα becomes phosphorylated, ubiquitinated and subsequently degraded by the proteasome. The de-inhibited p50:p65 heterodimers then enter the nucleus and regulate the expression of NF-κB target genes^12,13^. However, p50 and p52 may also form homodimers, presumably functioning as transcriptional repressors, as these homodimers lack a transactivation domain which is only provided by the Rel proteins.

*NFKB1* mutations are associated with a broad phenotype ranging from clinically unaffected to antibody deficiency and immune dysregulation within members of the same family^1,7,14^. Affected mutation carriers have a high frequency of autoimmunity, lymphoproliferation, noninfectious enteropathy, opportunistic infections, autoinflammation and malignancy^2,3,7^. Many patients have reduced numbers of class-switch memory B cells with an accumulation of CD21low B cells while maintaining an overall normal T-cell phenotype^15–17^.

The observation of *NFKB1* haploinsufficiency mutations suggests that *NFKB1*-related diseases result in *decreased* NF-κB signaling^18^ due to insufficient availability of signaling components. This observation is supported by reports of RelA haploinsufficiency^19,20^. Nevertheless, as shown by the existence of A20 deficiency, a negative regulator of the NF-κB signaling pathway, immune dysregulation can also be a result of an *increased* NF-κB activation^21–23^. We previously demonstrated that *NFKB1* mutations cluster into at least four different groups: N-terminal truncating mutations causing p50 haploinsufficiency, truncating mutations in the central part that predict immediate expression of p50-like proteins (precursor skipping), missense mutation affecting only the p105 precursor, and missense mutations affecting both, p105 and the active p50 (N-terminal half)^7,24^. An increase of p50 molecules by e.g. precursor-skipping mutations may support the formation of p50 homodimers, which would in turn repress gene transcription (as they lack the TAD domain of the Rel proteins) and therefore block NF-κB signaling. Clinical observations, however, taught us that *inhibition* of proinflammatory NF-κB signaling e.g. by anti-TNFα or anti–IL1 therapy ameliorated disease activity (Franxman et.al.^25^, Toh et.al.^26^, Dieli-Crimi et.al.^16^, and unpublished observation B. Grimbacher).

Thus, for a research evidence-based treatment of the patient, it is crucial to understand the detailed impact of the identified *NFKB1* variants on the NF-κB signaling processes, its chromatin regulation, the role of posttranscriptional modifications, and crosstalk between NF-κB and other signaling pathways, in the context of disease. Because of the nature of the genetic mutations in *NFKB1*, downstream effects on NF-κB target genes are expected. These NF-κB target genes, however, may lead to the expected pro-inflammatory response and regulate differentiation and effector function in lymphocytes^27,28^. Yet, there is also a substantial number of NF-κB target genes which are involved in autoregulatory negative feedback loops to limit NF-κB signaling^29^. Therefore, the assessment of expression profiles of these genes may provide us with a more complete and quantitative measure of the biological processes involved in disease expressivity and penetrance caused by the heterozygous mutations in *NFKB1*. Moreover, deeper mechanistic insights into the pathogenesis of NF-κB immune dysregulation are urgently needed to facilitate the identification of specific targets for therapy.

Recent progress in genomic and transcriptomic sequencing analysis elucidated the genetic landscape of several immune diseases^30–32^. These multi-omics strategies have already revealed new mechanisms and potential targets to develop novel therapies. Here, we performed an integrative genomic, epigenomic, transcriptomic, and proteomic analysis in B cells of *NFKB1* mutation carriers compared to individuals with wild-type *NFKB1* (WT). The resulting data serve as an important resource for future biological, diagnostic, and drug discovery efforts.

## Materials and methods

### Study Participants

In this study we included 13 individuals with heterozygous pathogenic mutations in *NFKB1* from six different families (six affected and seven unaffected *NFKB1* mutation carriers) and 23 unrelated control individuals tested WT for *NFKB1*. Written consent was obtained from all individuals. This study was approved by the institutional review board and conducted according to the ethical standards of the Helsinki Declaration. Approved ethics protocols are iPAD EC No.76/19, and for the NFKB Study protocol No. 295/13_200149 and 93/18_191111, from the University Hospital of the Albert-Ludwigs-University, Freiburg, Germany.

### Whole-exome sequencing

Genomic DNA was purified from human peripheral blood mononuclear cells (PBMCs) using QIAamp kits (Qiagen, Hilden, Germany) according to the manufacturer’s protocol. Whole-exome sequencing (WES) was performed using the custom SureSelect exome sequencing protocol from Agilent (Waldbronn, Germany). Exomes were enriched by using SureSelect exome v5 probes. Libraries were sequenced twice (two flow cells) on a HiSeq 2500 v4 with a 2 × 76 bp protocol generating four raw sequence data files (FASTQ) per sample. Data preprocessing was performed according to the GATK best practices and involved the following steps: (1) conversion of FASTQ files into an unmapped BAM file (PICARD tool FastqToSam), (2) addition of tags to the Illumina adapter sequences of the unmapped BAM file (PICARD tool MarkIlluminaAdapters), (3) conversion of the unmapped tagged BAM file into a FASTQ file (PICARD tool SamToFastq), (4) alignment to the reference genome build UCSC hg38 (BWA MEM), (5) identification of duplicated reads (MarkDuplicates PICARD), (6) BAM recalibration, and (7) indel realignment. Variant calling was performed with three different variant callers: GATK Haplotype caller, FreeBayes, and SAMtools. BASH and R scripts were subsequently used to (1) merge the VCF files, (2) identify and unify dinucleotide changes, and (3) format the data sets for importation into an inhouse specialized SQL database (GemmaDB) at the Center for Chronic Immunodeficiency in Freiburg. Variant annotation was performed using Ensembl’s Variant Effect Predictor tool (ensemble.org), and allele frequency (AF) data were extracted from the gnomAD exome (v2.1.1) and genome (v3) data sets (GnomAD browser). Individual frequencies were obtained by transforming the gnomAD AF data. Variant filtering was performed by selecting variants with (1) an individual frequency below 1% in both our internal cohort and the gnomAD (exomes or genomes) populations, which included control cohorts, such as those in the NHLBI-GO Exome Sequencing Project or the 1000 Genomes project, (2) a “high” or “moderate” predicted impact, (3) an alternative AF larger than 0.3 and read depth larger than 20, and (4) a zygosity matching an autosomal recessive or X-linked recessive mode of inheritance.

### Confirmation of Identified Sequence Variants by Sanger-Sequencing

DNA isolation was performed from blood samples treated with EDTA according to our local protocol using QIAamp kits (Qiagen, Hilden, Germany). *NFKB1* exons were amplified by PCR from gDNA and processed for Sanger sequencing according to standard protocols. Primer sequences are available in Supp.Table S1.

### Peripheral Blood Mononuclear Cells isolation and FACS sorting from isolated cells

Peripheral Blood Mononuclear Cells (PBMCs) were isolated from EDTA blood by Ficoll density centrifugation following standard protocols. Cells were centrifugated for 5 minutes, 300g at RT, and the supernatant was discarded. The cell pellet was resuspended with 100µl of the corresponding master mix containing optimized dilutions of antibodies, for 30 minutes at 4°C. Next, cells were washed with FACS buffer and centrifuged for 5 minutes, 300g at RT. The supernatant was discarded and the cell pellet was resuspended in 200µl of FACS buffer for sorting.

In a first approach, 10 patient and 19 WT PBMC samples were sorted into naïve B cells (CD19+IgD, CD27-) and stimulated for 24 hours. B cell sorting was conducted with: CD19 APC-Cy7, CD20 APC, CD21 PE-Cy7, CD27 Brilliant Violet 421, CD38 PerCP-Cy5.5, IgM Alexa Fluor 488, IgD PE. Cells were sorted on a MoFlo Astrios Cell sorter (Beckman Coulter, Brea, CA, USA). Naïve B lymphocytes (250.000 cells/ml) were seeded on a 96 well round bottom plate in RPMI medium supplemented with 10% FCS and 1% Penicillin-Streptomycin. These cells were stimulated with 1µg/ml CD40L (provided by Pascal Schneider, Lausanne, Switzerland) and 50ng/ml IL21 (Miltenyi) and incubated for 24 hours at 37ºC, 5% CO2. After stimulation, cell numbers were determined with a Neubauer chamber. A total of 5,000-50,000 cells were used for DNA and RNA extraction, 5,000-50,000 were used for ATAC-sequencing and 74,000-200,000 for proteome analysis. B cell activation (CD69 APC, CD80 Brilliant Violet 421, CD86 PerCP-Cy5.5, CD95 Brilliant Violet 650, HLA-DR PE-Cy7) was measured on a B LSR Fortessa from Becton Dickinson (BD Biosciences, San Jose,CA).

On a second approach, PBMCs from three patients with pathogenic *NFKB1* variants and four WT control individuals were sorted into four different B cell subtypes: naïve (CD19+, CD21+, IgM+, IgD+, CD27, CD38-), IgM+ memory (CD19+, CD21+, IgM+, IgD, CD27+, CD38-), switched memory (CD19+, CD21+, IgM-, IgD-,CD27+,CD38-) and CD21low (CD19+,CD21-, IgM+/-, IgD+/-, CD27+/-, CD38-). B cell sort: IgM Alexa Flour 488, IgD PE, CD38 PerCP-Cy5.5, CD21 PE-Cy7, CD19 APC, CD27 BV450. Cells were sorted with the Cell sorter (Beckman Coulter, Brea, CA, USA). The cell pellets were subjected to DNA and RNA isolation.

### RNA / DNA isolation and sequencing

DNA and RNA were extracted from isolated cells using a Qiagen AllPrep DNA/RNA Micro Kit. DNA concentration was determined using a Qubit fluorometer (Termo Fisher Scientific). Library preparation and sequencing by Reduced Represented Bisulfite Sequencing (RRBS) was performed at the CeMM Research Center for Molecular Medicine of the Austrian Academy of Sciences in Vienna, Austria, according to an established and extensively validated protocol^33–35^.

The RNA library preparation and the sequencing was performed by Novogene (Cambridge, UK). The NEBNext RNA First Strand Synthesis Module and NEBNext Ultra Directional Second Strand Synthesis Module were used for low input RNA samples. The low input RNA concentrations from different healthy controls were sequenced using different library preparations and they were tested for the best sequencing outcome. The tested library preparations were the above mentioned low-input protocol, an rRNA depletion protocol and a SMARTer amplification protocol. After quality control, it was decided to process all samples from patients with the SMARTer amplification protocol, due to the higher number of unique reads. Samples were sequenced using the NovaSeq 6000 (Illumina) in paired-end mode. Generated reads had a length of 150 bp.

### DNA methylation analysis

DNA was isolated from 5,000-50,000 FACS purified cells using the Qiagen DNA/RNA Micro Kit. RnBeads v2.12.2 was used for the analysis with standard settings^36^. Except for the addition of a set annotation for regulatory elements taken from Ensembl Regulatory Bild from BLUEPRINT data, hg38 assembly. Specifically, transcription factor binding sites, CTCF-binding sites, proximal and distal enhancers were used to study their overlap with differentially methylated regions using a custom RnBeads workflow. DNA Methylation values were calculated across 500bp regions. Only windows that contained ≥5 CpG sites were considered. The single nearest gene option was used for the association between the different genomic regions and genes.

### RNA data analysis

RNA was isolated from 5,000-50,000 FACS-purified cells using Qiagen AllPrep DNA/RNA Micro Kit. The fastq files were uploaded to the Galaxy web platform and the public server usegalaxy.eu was used for the first part of the analysis ^37^. Quality control of raw reads was done using FastQC. Quality Trimming was performed with TrimGalore, 10bp were removed from the 5’ end of each read. Sequences were aligned to the reference genome (GRCh38/hg38) using RNA-STAR. Default parameters for paired-end were used, except for the modification of the length of the genomic sequence around annotated junctions to 149 and the maximum number of alignments to output a read’s alignment results to 3. Duplicates were not removed. The gene expression was measured with FeatureCounts using built-in gene annotations, unstranded specification and default parameters. Further analysis was performed in R v4.2.0 with an R custom script. Genes with read counts lower than 100 were excluded from the analysis. Differentially expressed gene analysis was done using R Bioconductor package DESeq2 v.3.15. Batch effect was corrected. Variance stabilizing transformation was applied to DESeq2 results. Differential expressed genes were filtered according to their adjusted p-value (<0.05) and absolute value of log2 fold change (>0.5). Principal component analysis was carried out after batch correction using Bioconductor limma package. Heatmaps and volcano plots were generated using the R package pheatmap, and the Bioconductor package EnhancedVolcano, respectively. Gene set enrichment analysis (GSEA) was performed using the R Bioconductor package fgsea together with the Hallmark, and KEGG pathways from the M SigDb. All differentially expressed genes were used for GSEA. Ggplot2 package was used to create dot plots of pathway analysis.

### Protein extraction and enzymatic digestion

Between 74.000-200.000 FACS-sorted cells were washed twice in phosphate-buffered saline (PBS), and dry pellets were flash frozen and stored at -80°C. Further processing was performed in the Institute Biochemistry in Bellinzona., Switzerland. Cell pellets were lysed in 50 µl of 8M urea in 50 mM ammonium bicarbonate (ABC) followed by sonication (15 minutes at 4°C), with a Bioruptor (Diagenode). Up to 100µg of protein extract per sample was reduced with 10 mM dithiothreitol (20 minutes at RT). Subsequently, proteins from each sample were alkylated with 50mM iodoacetamide (30 minutes at RT) and digested with 1 µg of LysC (FUJIFILM Wako) in 8M urea, 50 mM ABC (2 hours at RT). After digestion, the buffer was diluted with 50 mM ABC to a final of 2M urea and digested overnight at RT with 1 µg trypsin (Promega, Walldorf, Germany). On the next day, the addition of acetonitrile (ACN) to 2% and trifluoroacetic acid (TFA) to 0.3% terminated the trypsin digestion. The samples were centrifuged 5 minutes at maximum speed, and the supernatants were desalted using C18 StageTips. Peptide elution was achieved with 80% ACN, 0.5% acetic acid. To evaporate the solution, vacuum centrifugation was performed followed by resuspension of the purified peptides with 2% ACN, 0.5% acetic acid, 0.1% TFA. Up to 1 µg of purified peptides per sample were injected in a liquid chromatograph-tandem mass spectrometer, as single-shot measurements.

### Liquid chromatography– with tandem mass spectrometry (LC-MS/MS) analysis

Peptide separation in buffer A (0.1% formic acid) was achieved using an EASY-nLC 1200 HPLC system (Thermo Fisher Scientific) and a column (75 µm inner diameter, 50 cm length, ReproSil-Pur C18-AQ 1.9 µm). Elution was carried out in a 5-30% buffer B (80% ACN, 0.1% formic acid) linear gradient for 150 minutes and a flow rate of 250 nl/min. Ions were produced by a nanoelectrospray source (Thermo Fisher Scientific) and fragments feed into a Q Exactive HF mass spectrometer (Thermo Fisher Scientific). Measurements were performed in a data-dependent mode. Parameter settings in Xcalibur software (Thermo Scientific) comprised a survey scan range of 300-1,650 m/z, resolution of 60,000 at 200 m/z, maximum injection time of 20 ms and AGC target of 3e6. In a 1.7 m/z isolation window, up to ten ions with high frequency and a charge of 2-5 were isolated. Next, the ions underwent higher-energy collisional dissociation (HCD) fragmentation with a normalized collision energy of 27 to acquire MS/MS scans. Repeated sequencing was excluded by setting dynamic exclusion to 30 s. Measurement parameters comprised a resolution of 15,000 at 200 m/z, maximum injection time of 55 ms and AGC target of 1e5.

### ATAC-seq processing

Accessible chromatin mapping was performed using the ATAC-seq method as previously described^38^ with parameters reported in Ramirez et al^39^. In brief, 5,000-50.000 cells per experiment were centrifuged 5 minutes at 8°C, and resuspended in 25 µl transposase mixture (12.5 µl Tagmentation DNA buffer, 1 µl Tn5 transposase (Illumina), 10.75 µl nuclease-free water, 0.25 µl 1% digitonin (Promega) and 0.5 µl 50x Protease Inhibitor cocktail (Roche)). After incubation of the solution, 30 minutes at 37°C, and DNA purification, the DNA was eluted in 13.5 µl 10mM Tris-HCI, pH 8.5. Further library preparation (NEBNext Ultra II kit) and sequencing, 50bp single-end on a HiSeq 4000 was performed at the CeMM Research Center for Molecular Medicine of the Austrian Academy of Sciences in Vienna, Austria.

### ATAC-seq analysis

Unaligned reads were quality controlled using FastQC. Reads were then trimmed, aligned to hg38, and filtered using Skewer, Bowtie2^40^, and Sambamba^41^ respectively. As previously described, all downstream analyses were performed on the filtered reads^42^ (>100 million). Reads mapping to mitochondrial DNA were excluded before peak calling was performed using HOMER^43^. The parameters ‘-style factor and -L 20’ were used to analyze transcription factors (20-fold greater tag density than in the surrounding 10kb region) and ‘-style super’ was used to analyze super enhancers with default settings. With ‘getDifferentialPeaksreplicates.pl -balanced –edgeR -L 10’ parameters (10-fold greater tag density than background), differential transcription factor peaks were identified. Peaks were merged for the same cell type using mergePeaks and annotated with annotatePeaks.pl. To perform principal component analysis, a non-annotated file generated with the ‘noann’ option was uploaded to Rstudio and visualized with ggplot2 (version 3.3.5). Transcription factor binding motifs were identified with HOMER function findMotifsGenome.pl and ‘-size 200 (peak size) and -len 8,10,12,15 (motif length)’ settings. Next, a matrix file was generated using HOMER with ‘-size 200 -hist 400 (bin size in bp) -ghist (outputs gene profiles for each gene) parameters. The matrix file was used for unsupervised hierarchical clustering using Cluster 3.0 (parameters: normalize genes, cluster genes and array, similarity metric: correlated uncentered) and visualized with Java Tree View.

### Proteome data analysis

Using the MaxQuant software, v.1.6.7.0^44^ and the integrated Andromeda search engine^45^, peptides and proteins with a false discovery rate of < 1% were determined from the Xcalibur raw files. The Human UniProt database (June 2019) and a common contaminants database were used for peptide and protein identification with “Trypsin/P” enzyme specificity, a maximum of 2 missed cleavages and minimum length of 7. Variable modifications included N-terminal protein acetylation and methionine oxidation, whereas cysteine carbamidomethylation was selected as fixed modification. To compare runs among each other, mass and normalized retention time (time window of 0.7 minutes and an alignment time window of 20 minutes) were used. The MaxLFQ algorithm^46^ was used to quantify label-free proteins (LFQ) with a minimum peptide ratio count of 1 and LFQ intensities were log2 transformed. Proteins only identified by site, reverse peptides, and potential contaminants were removed. In Rstudio, the R Bioconductor package DEP (version 1.12.0) was used for normalization and imputation with a left-censored method (0.3 width and 1.8 downshift). The t-test from the R package limma (version 3.46.0) was used to determine differently expressed proteins (alpha = 0.05, absolute log2 fold change = 1.5). All visualizations were performed as described for the transcriptome analysis.

### Association between DNA methylation, open chromatin and Gene Expression

For association analysis, samples that had DNA methylation, ATAC-seq and/or gene expression data were examined. The Bioconductor package Complex Heatmap (v2.10.0) with a custom R script were used for the association analysis between gene expression, DNA methylation, and ATAC-seq. For methylation and ATAC-seq data, methylation and preferentially open chromatin regions from *NFKB1* mutation carriers were used for the analysis. For RNA-seq and proteome data, differential gene or protein expression between *NFKB1* mutation carriers and WT controls were used. The median methylation is represented as a beta-value from 0 to 1 (0 to 100% methylation). For ATAC-seq data, position adjusted reads from initial peak regions were used. For differentially expressed genes in the RNA-seq, data log2fold changes values were used, as were for differentially expressed proteins. Venn diagrams were drawn using the R package ggVennDiagram. We performed a supervised analysis of NF-κB target genes on B cells in all the data sets. The list of NFκB -target genes was adapted from https://www.bu.edu/nf-kb/gene-resources/target-genes/, and an additional search with PubMed.

## Results

### Clinical and genetic characterization of patients

In this study, we included a total of 13 *NFKB1* mutation carriers from six different families and 23 individuals WT for *NFKB1* (Table 1). Among the mutation carriers, six were classified as affected and seven were unaffected. All affected individuals presented with hypogammaglobulinemia and a history of recurrent respiratory infections. The median age of mutation carriers was 55.5 (±21.28) years. The clinical phenotype was characterized by lymphoproliferation, particularly splenomegaly and lymphadenopathy. Autoimmunity and inflammatory gastrointestinal involvement were present in all affected individuals. At the time of analysis, all but one affected mutation carrier (FR097.2) received immunomodulatory treatment (Table 2).

**Table 1.**
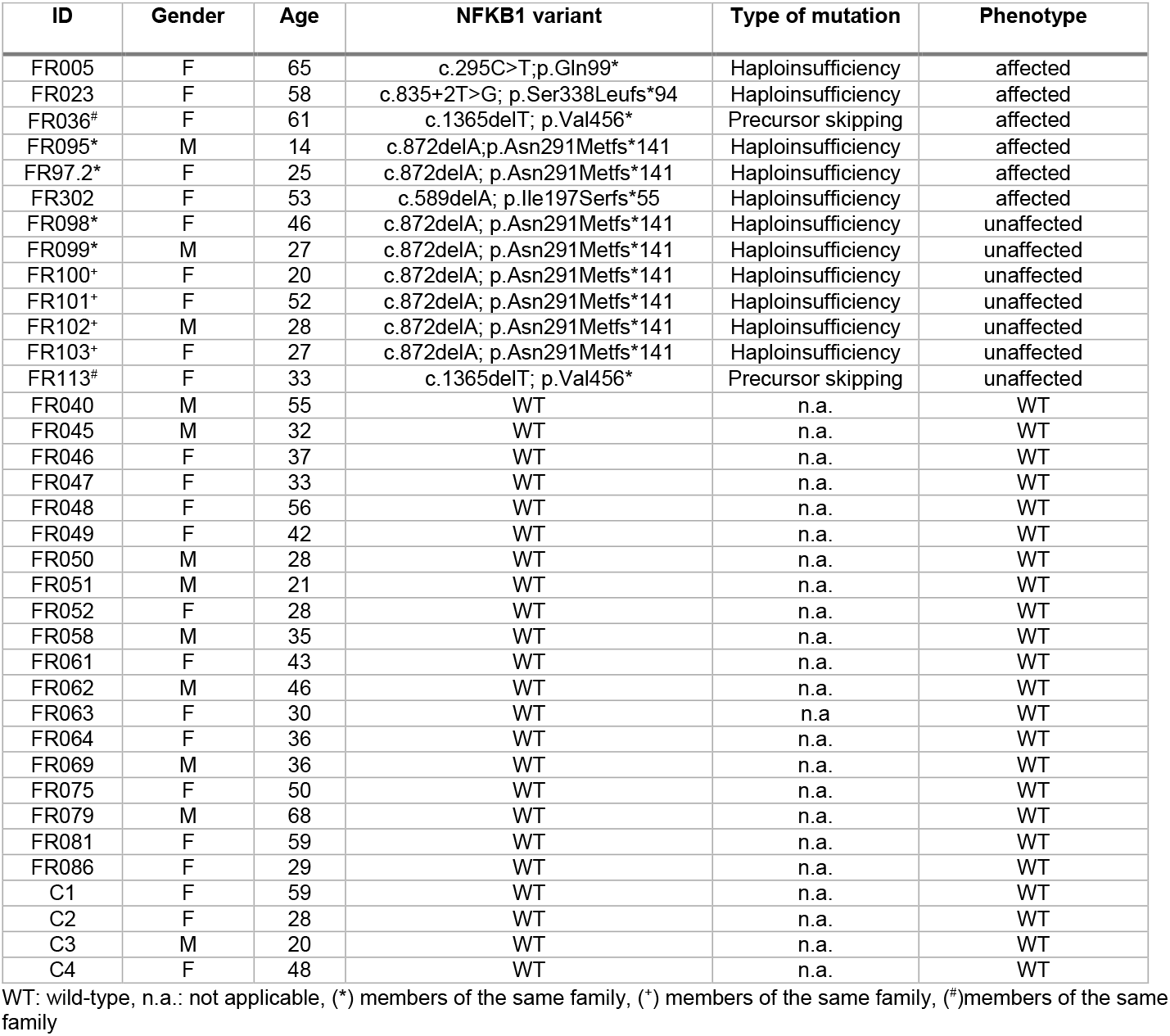
List of individuals included in this study

**Table 2.** Clinical characteristics of affected mutation carriers

| ID | Gender | Age | NFKB1 mutation | Category | Respiratory involvement | Gastrointestinal involvement | Lymphoproliferation | Autoimmunity | Others | Immunomodulation |
| --- | --- | --- | --- | --- | --- | --- | --- | --- | --- | --- |
| FR005 | F | 65 | c.295C>T;p.Gln99* | Haploinsufficiency | URTI, asthma | Celiac like disease, hepatopathy | Splenomegaly, lymphadenopathy | Enteropathy | Osteoporosis, Norovirus infection, cold thyroid nodules | Glucocorticoids |
| FR023 | F | 58 | c.835+2T>G;p.Ser338Leufs*94 | Haploinsufficiency | URTI, GLILD | Liver cirrhosis | Splenomegaly | AITP, AIHA, Arthritis, CNS vasculitis | Osteopenia | Glucocorticoids, Upadacitinib |
| FR036 | F | 61 | c.1365delT; p.Val456* | Precursor skipping | URTI, GLILD | CMV infection, hepatitis | Splenomegaly, lymphadenopathy | AITP, AIHA, enteropathy, arthritis | no | Glucocorticoids, Abatacept |
| FR095* | M | 14 | c.872delA; p.Asn291Metfs*141 | Haploinsufficiency | URTI, pneumonia | IBD-like disease | no | Autoimmune thyroiditis, arthritis | Apthous ulcerations | Glucocorticoids, Abatacept |
| FR97.2* | F | 25 | c.872delA; p.Asn291Metfs*141 | Haploinsufficiency | URTI | IBD-like disease | no | Arthritis | no | no |
| FR302 | F | 53 | c.589delA; p.Ile197Serfs*55 | Haploinsufficiency | URTI, pneumonia | Chronic diarrhea | Splenomegaly, lymphadenopathy | Enteropathy | no | Glucocorticoids |

URTI: upper respiratory infections, GLILD: Granulomatous lymphocytic interstitial lung disease, CMV: cytomegalovirus, AITP: autoimmune thrombocytopenia, AIHA: autoimmune hemolytic anemia, CNS: central nervous system, IBD: inflammatory bowel disease, HCV, (\*) members of the same family

All *NFKB1* mutation carriers underwent WES for mutation detection and Sanger sequencing to verify the presence of the variant. All variants with a frequency of less than 1% in gnomAD (https://gnomad.broadinstitute.org) which were detected in our patients by WES are displayed in the supplementary Table S2. Eleven individuals carried N-terminal truncating mutations, predicted to cause haploinsufficiency of the p105 precursor protein, and the mature p50. Patient FR005 carried the mutation c.295C>T; p.Gln99*, it was previously described ^5^. The single base pair deletion (c.872delA; p.Asn291Metfs*141) was present in eight patients (from two different families). Patients FR095 and FR097.2 are affected and members of the same family as patients FR098 and FR099. In the other family, all patients (FR100, FR101, FR102 and FR103) were unaffected. All individuals were previously reported^78^. Patients FR023 and FR302 carry the haploinsufficiency mutations c.835+2T>G (p.Ser338Leufs*94) and c.589delA (p.Ile197Serfs*55), respectively. The mutation of patient FR023 was previously reported^7^. In addition, two patients, including one affected (FR036), and one unaffected (FR113) member from the same family carried a precursor skipping mutation c.1365delT (p.Val456*). These types of mutations affect the central part of p105 and predict the immediate expression of a p50-like protein thereby skipping the precursor stage. However, all eleven patients with pathogenic mutations in *NFKB1* also carried rare variants (<1% in gnomAD) in NF-κB downstream target genes (Table 3).

**Table 3.**
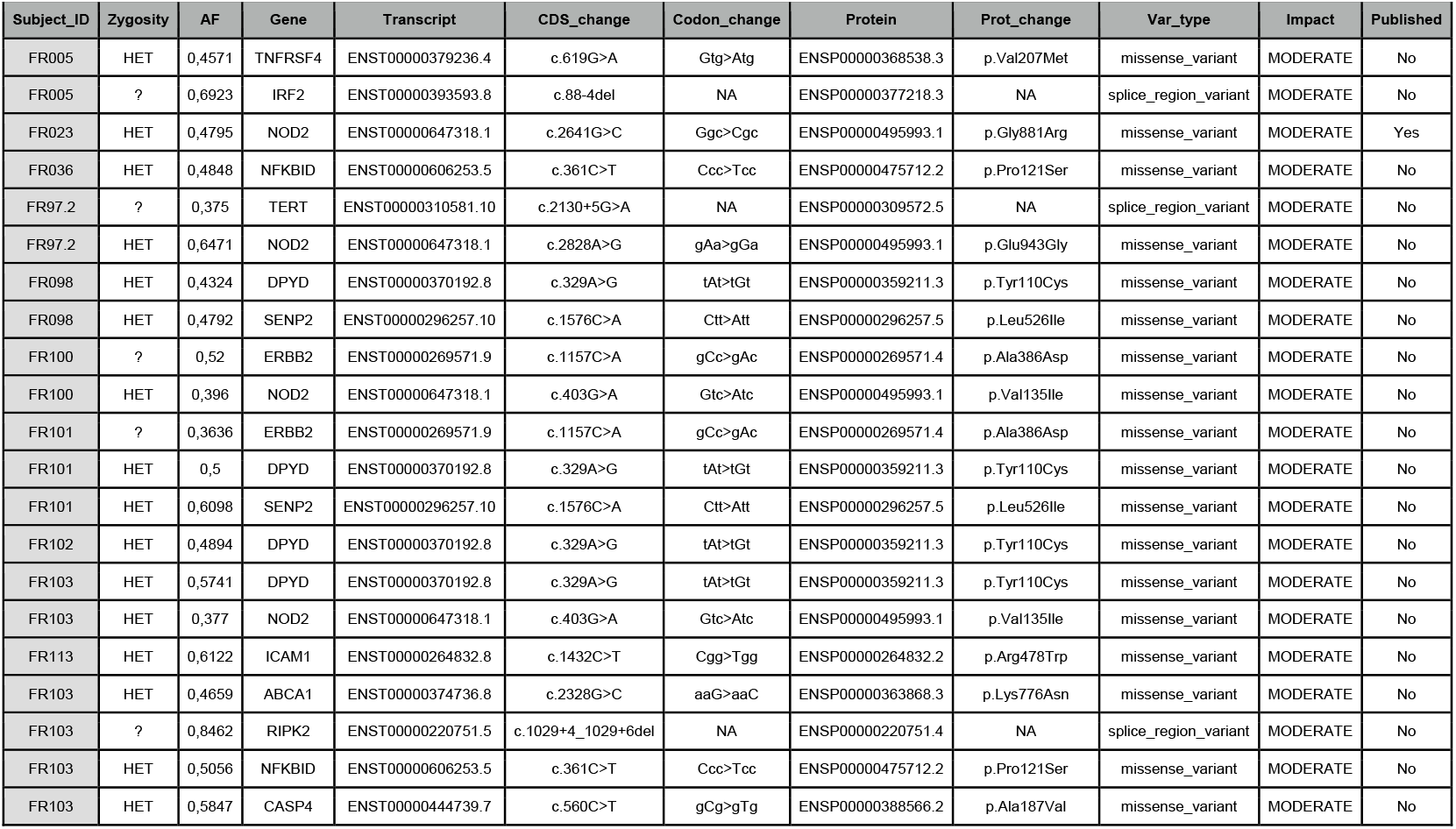
List of Rare variants (<1% in gnomAD) in NF-κB downstream target genes of *NFKB1* mutation carriers

### Abnormal expression of NF-κB target genes in unstimulated and stimulated B cell subsets indicates reduced expression of negative regulators

Heterodimeric NF-κB transcription factors (composed of p50 and one of the Rel proteins) act as inducers of gene expression^47,48^. We determined the gene expression profiles of both sorted unstimulated naïve B cells, and CD40L/IL21-stimulated naïve B cells derived from *NFKB1* mutation carriers by RNA seq. We obtained RNA sequencing data of stimulated naïve B cells from ten *NFKB1* mutation carriers (three affected and seven unaffected), and 18 age- and gender-matched healthy donors, WT for *NFKB1*. In addition, we performed RNA sequencing on four unstimulated B cell subtypes (naïve-, IgM memory, switched memory, and CD21low-B cells) from three affected *NFKB1* mutation carriers (FR095, FR103, and FR302) and 4 WT. A list of sequencing data available per sample is listed in Supp. Table S3.

Since IL21 and CD40L stimulation induce a strong activation of NF-κB signaling, leading B cells to a proliferative burst and later plasma cell differentiation^49^, we inspected the transcriptome of 24h activated naïve B cells on ten *NFKB1* mutation carriers (three affected and seven unaffected) and 18 NFKB1 WT individuals. In comparison to wildtype controls, stimulated naïve B cells from NFKB1 mutation carriers displayed profound changes in NF-κB target gene expression. This observation suggests that the pathogenic effect of these variants is evident at the transcriptional level (column 1, Figure 1a). Particularly, the increased expression of *NR4A2* and *GZMB*, in combination with the decreased mRNA expression of *TNFAIP3, TRAF1, TRAF6, TNIP1 and CYLD*, points towards an induction of B cell proliferation and dysregulation of NF-κB inhibitors^18^. Interestingly, some B cells secrete granzyme B (codified by *GZMB*) when recognizing viral antigens, in the setting of infection, in autoimmune diseases, in chronic lymphocytic B cell leukemia, and when infiltrating solid tumors ^50–52^. Therefore, *NFKB1* haploinsufficiency seems to mimic this state of a chronic stimulus such as present in chronic virally infected B cells.

**Fig 1.**
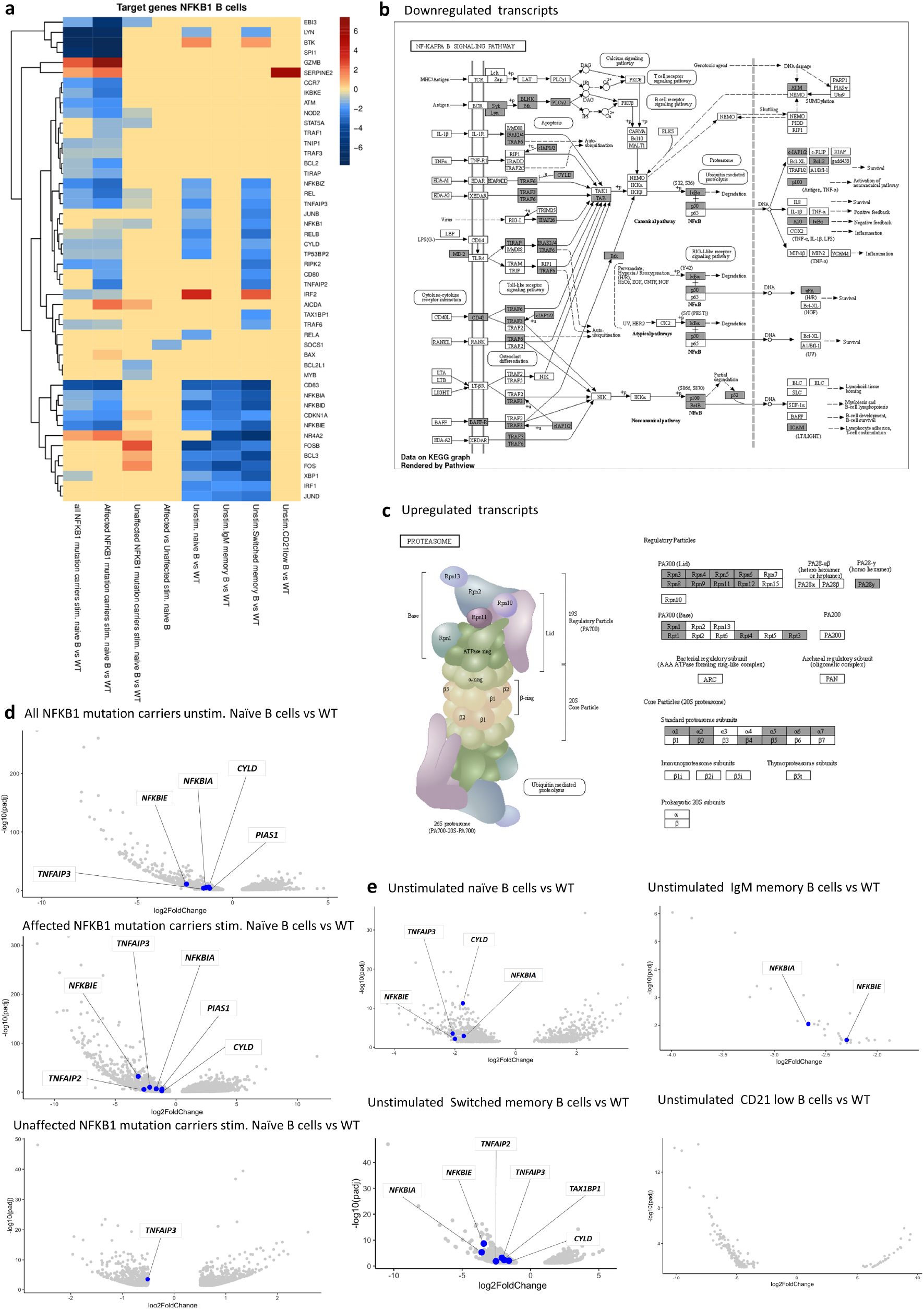
Differential gene expression pattern in B cell subsets derived from NFKB1 mutation carriers indicates impairment of NFKB negative regulators and NFKB target genes. **a)** Heatmap representation of differentially expressed NFKB target genes in WT and NFKB1 mutation carriers (log2 fold change ≥0.5, and adjusted p-value <0.5). Columns show the comparisons on unstimulated naïve-, stimulated naïve-, IgM memory-, and switched memory-B cells between NFKB1-mutated and WT. Color represents genes that are differentially expressed, where blue and red represent a decreased and an increased expression, respectively. **b)** Pathway overview of decreased expression of NFKB target genes comparing NFKB1 affected mutation carriers and WT. Differentially expressed genes are statically significant, with an adjusted p-value <0,05, and are highlighted in gray. Figure shows decrease expression of negative regulators of the NF-κB signaling pathway (*CYLD, TNP1, TRAF1, TNFAPI2 and TNFAIP3* (A20)) in affected NFKB1 mutation carriers on stimulated B cells. **c)** KEGG pathway representation of NFKB target genes with increased expression on stimulated naïve B cells in affected NFKB1 mutation carriers compared to WT. Gray color indicates statically significant differentially expressed genes (adjusted p-value <0,05). Genes encoding proteasome subunits showed elevated mRNA expression. **d)** Volcano plot representation of the NFKB target genes transcriptomic comparison of affected plus unaffected-, affected- and unaffected-NFKB1 mutation carriers with WT. In blue, reduced expressed genes that negative regulate NF-kB signaling pathway on stimulated naïve B cells (log2 fold change ≤0.5, and adjusted p-value <0.5). **e)** Volcano plot representation of NFKB target genes at the transcriptomic level. NFKB1 mutation carriers were compared to WT across different unstimulated B cell subsets. Plots show data from unstimulated naïve-, IgM memory-, switched memory-, and CD21 low-B cells. Blue color dots represent reduced expressed genes with a role in the negative regulation of NF-kB signaling pathway (log2 fold change ≤0.5, and adjusted p-value <0.5). There were no differentially expressed negative regulator genes of the NFKB signaling pathway on CD21-lowB cells compared to WT.

We then asked whether the disease status of each patient might influence NF-κB target gene expression i.e. whether affected and unaffected mutation carriers display unique NF-κB target gene expression profiles. Thus, we split the analysis of the ten stimulated naïve B cells *NFKB1*-mutated samples into two groups. The first group comprised three patients, who were clinically affected at the time of blood drawing. The second group contained seven unaffected mutation carriers. Data analysis presented in columns 2 and 3 in Figure 1a shows that the affected *NFKB1* mutation carriers accounted for the strong NF-κB target gene dysregulation when compared to matched healthy controls, while the seven unaffected *NFKB1* mutation carriers remained with almost normal levels of NF-κB target gene expression. When comparing affected *versus* unaffected *NFKB1* mutation carriers, *SOCS1* had lower expression levels (column 4, Fig 1a.). *SOCS1* is one component of the ubiquitin ligase complex that mediates NF-κB degradation^53^, suggesting that the ubiquitination process of NF-κB subunits in the control of transcription is impaired.

In summary, columns 1-4 in Figure 1a show that stimulated naïve B cells of *NFKB1* mutation carriers have a significantly different gene expression pattern when compared to WT. To investigate which biological processes are most affected, we performed pathway analysis utilizing hallmark gene sets. In *NFKB1* mutation carriers, we found TNF-α signaling *via* NF-κB, and the interferon gamma (IFN-γ) response to be under-expressed.

Next, we compared the data sets of unstimulated naïve B cells from three NFKB1 mutation carriers (FR095, FR103, and FR302) to four matching healthy control samples. We observed that unstimulated naïve B cells from *NFKB1* mutation carriers showed changes of NF-κB target gene expression (column 5, Figure 1a) with overall lower expression levels compared to WT. As expected, under resting conditions, the haploinsufficiency mutation leads to a reduced production of p105, hence also a reduced amount of its processed form p50, and therefore to disturbances of the baseline expression of NF-κB target genes^1^.

We were also interested whether other unstimulated B cell subsets such as IgM memory-, switched memory-, or CD21-low B cells showed a dysregulation of NF-κB target genes. Whereas CD21-low B cells showed only minor variation when compared to healthy donors, we found clearly reduced expression levels for a distinguished set of NF-κB target genes in IgM memory- and switched memory B cell compartments. For instance, genes codifying for NF-κB inhibitors *NFKBIA, NFKBID, NFKBIE*, and transcription factors essential during B cell development (*BCL3* and *XBP1*) were strongly dysregulated (columns 6-8, Figure 1a). This observation goes along with the observation that NFKB1 patients present with impaired terminal B cell differentiation.

To test how *NFKB1* mutations affect downstream pathways, we performed a KEGG enrichment analysis on differentially expressed NF-κB-target genes. We compared stimulated naïve B cells from affected mutation carriers (three affected) to the WT control group. In patient samples, we found reduced expression levels of genes of the NF-κB signaling pathway such as *ATM, BCL2, BIRC2 (c-IAP1), BTK, CD40, CYLD, LYN, NFKBIA(I*κBα), *PLAU (uPA), PLCG2, RELB, SYK, TAB, TIRAP, TRAF2, TRAF3*, and *TRAF6*. Importantly, expression of several negative regulators of the NF-κB signaling pathway was found to be reduced including *CYLD, TNP1, TRAF1, TNFAIP2 and TNFAIP3* (A20) (Figure 1b). Moreover, increased expression of core and regulatory components of the proteasome (20S and PA700) was observed (Figure 1c). The proteasome has an important role in the strict control of NF-κB activity. On one side, the ubiquitination and proteasomal degradation of IκB promotes NF-κB activity. On the other side, ubiquitination leads to degradation of NF-κB subunits and culminates its transcriptional activity^54^.

Next, we compared stimulated B cells derived from the entire group of *NFKB1* mutation carriers (n=10) with the WT control group. Consistent with the previous observations, we found a decreased expression of genes encoding for components of the NF-κB signaling pathway, like *BIRC2 (c-IAP1), BTK, CD40, CYLD, LYN, PLAU (uPA), PLCG2, RELB, TIRAP, TRAF3*, and *TRAF6* (Supp. Fig. S2), while regulatory components of the proteasome showed increased expression (Supp. Figure S3). In stimulated naïve B cells from unaffected *NFKB1* mutation carriers, we found reduced expression of both, genes of the NF-κB signaling pathway (*NFKB1* itself, *BCL2L1*, and *TNFAIP3*), and genes related to the TNF-α signaling pathway (*NOD2*, and again *NFKB1* and *TNFAIP3*). Some genes on the TNF-α signaling pathway had increased expression (AP-1, FOS, and BCL3), data shown in Supplementary Fig.S4-S6. From these results, we conclude that NF-κB signaling is active in mutated B cells, with reduced expression of several negative regulators of the pathway.

To achieve the termination of the NF-κB response the cell employs several mechanisms, including a negative feedback loop^24,29,55^. In the stimulated data set we observed a low expression of negative regulators of NF-κB in NFKB1 mutation carriers (Figure 1d). Remarkably, we also found lower expression of negative regulators of NF-κB in unstimulated naïve-, IgM memory-, and switched memory-B cells from *NFKB1* patients, but not in CD21low B cells (Fig. 1e). This was not surprising, given that CD21low B cells usually show a divergent RNA-seq profile^56^. This population has been associated with autoimmunity and chronic parasitic or viral infections^17^.

Our findings demonstrate that the transcriptional program of stimulated and unstimulated B cells across different subtypes is compromised in *NFKB1*-mutated cells. The lack of negative regulation may lead to chronic inflammation as observed in our patients, and in patients with rheumatoid arthritis^57^. Consistent with the transcriptional profile of stimulated naïve B cells, the limited expression of negative regulators of NF-κB signaling might explain the inflammatory phenotype of *NFKB1*-mutated individuals as well as the immune dysregulation. From these observations, we conclude that transcriptional changes are associated with clinical manifestations of immune dysregulation.

### Defective B cell differentiation and proliferation is caused by impairment of Chromatin accessibility in NFKB1-mutated cells

To study the enhancer landscapes in *NFKB1* mutation carriers, isolated B cells from three affected (FR005, FR023, and FR036) and five unaffected (FR099, FR100, FR101, FR102 and, FR103) *NFKB1* mutation carriers, and 17 wildtype control individuals were stimulated with CD40L/IL21 for 24h and analyzed for chromatin accessibility by ATAC-seq. 549 differentially accessible regions (DARs) were detected comparing *NFKB1* mutation carriers to WT. The majority of DARs were accessible in WT (381), while 168 DARs were accessible in mutation carriers (Figure 2a). To identify transcription factors associated with differential chromatin accessibility, we performed *de novo* transcription factor binding motif (TFBM) analyses. The *CTCF* motif was identified as the most enriched TFBM in *NFKB1* mutation carriers compared to WT, suggesting that the chromatin architecture is impaired. Furthermore, binding motifs for *NFAT, NKX2, NR1A2* and *STAT5* were found enriched, influencing B cell differentiation and proliferation. Enrichment in DNA-binding factors involved in cell activation and survival, such as *REL, SPIB, MZF1, ETV6* and *IRF-BATF*, were found enriched in WT individuals (Figure 2b). In order to identify a dysregulated chromatin landscape related to affected disease status, affected *NFKB1* mutation carriers were compared to WT. Consistent with the previous findings, one third of DARs (479) were accessible in affected *NFKB1* mutation carriers and two third of DARs (917) were accessible in WT (Fig. 2c). Similar TFBM were detected in WT, such as *SPIB, REL, IRF and BATF*. For *NFKB1* mutation carriers, *CTCF* was the motif with the highest enrichment, as in the previous analyses. CCCTC-binding factor (CTCF) is a chromatin architectural protein, which facilitates the interaction of different chromatin regions. It can thereby function as a repressor, transcriptional activator or insulator protein, and it can also recruit other transcription factors while bound to chromatin domain boundaries^58^. Furthermore, motifs for TFs involved in germinal center (GC) response were enriched such as *SP1, SOX4, IRF1*, and *PAX5* (Fig. 2d). In contrast to the affected *NFKB1* mutation carriers, the unaffected carriers presented with two third of DARs (248), while the WT had 129 DARs accessible (Fig. 2e). CTCF was the most enriched TFBM, as in previous analyses. The binding motif for STAT5 was detected, which was similarly enriched in the combined analysis of *NFKB1* mutation carriers. Transcription factors influencing B cell activation and development, such as *IKZF1* and *FRA1*, showed enriched motif accessibility. As observed in the previous analyses, *REL* and *SPIB* were enriched in WT (Fig. 2f). In addition, enrichment of the *NFKB2* binding motif was detected. Due to opposing results from affected and unaffected *NFKB1* mutation carriers, the two groups were compared to identify disease-specific signatures. Similar as in the comparison of unaffected *NFKB1* mutation carriers *vs* WT, two thirds of DARs (242) were accessible in the unaffected individuals, while one third of DARs (150) were accessible in affected *NFKB1* mutation carriers (Fig. 2g). *SIX2, PRDM4, OSR2*, Pu.1-IRF and *IRF5* were the most enriched TFBM in affected *NFKB1* mutation carriers. For unaffected *NFKB1* mutation carriers *SPI1, REL, AP-1, SOX10* and *SPIC* were most enriched TFBMs (Fig. 2h). With our identification of accessible chromatin and TFBM, our data reveal dysregulation of key players of B cell differentiation and proliferation. NF-κB (*REL*) and ETS family members (*SPIB, SPI1, SPIC*) seem most affected, as they were observed in all comparisons.

**Fig. 2.**
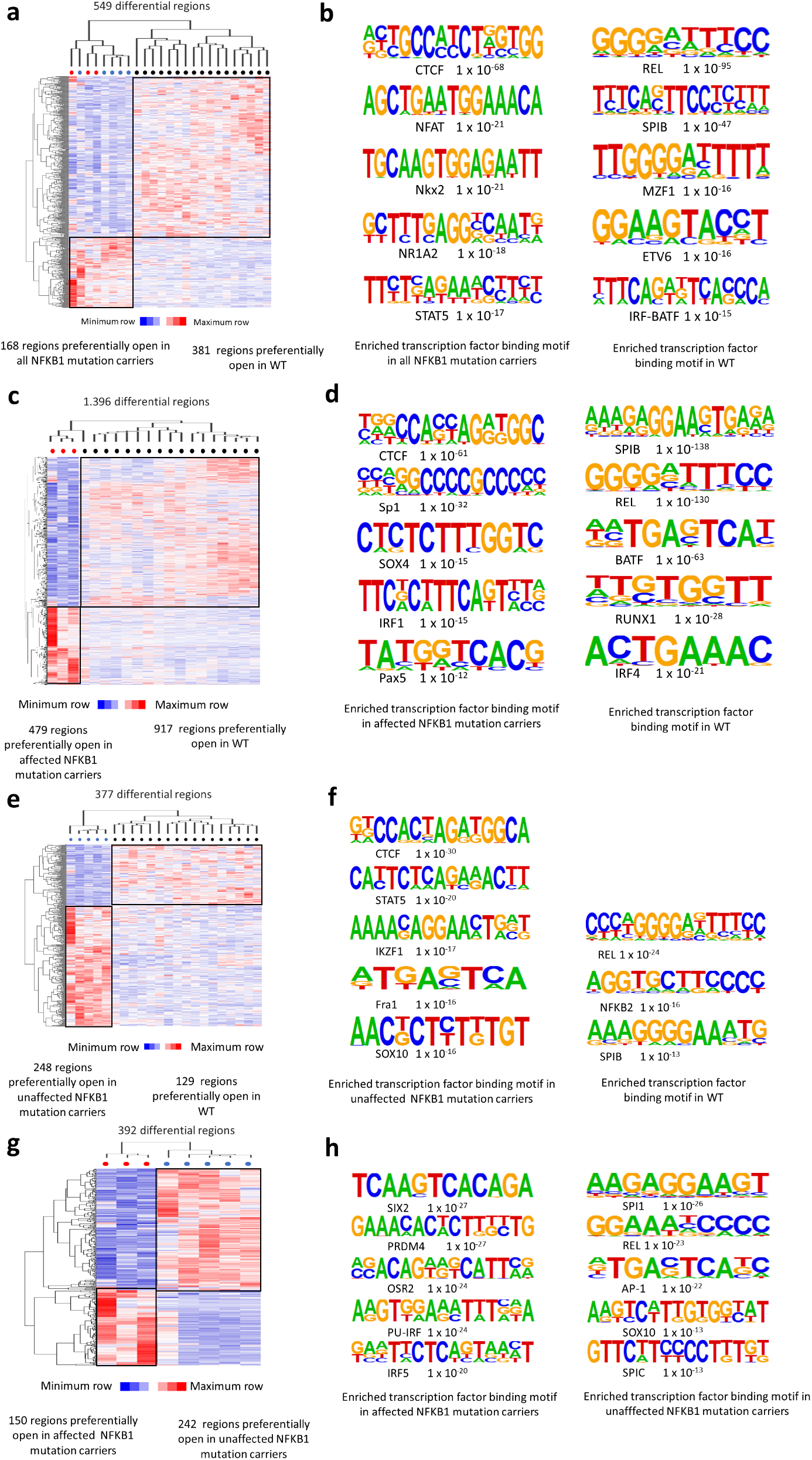
Chromatin accessibility defects in NFKB1 immune Dysregulation on stimulated naïve B cells. **a)** Clustered heatmap for the differentially accessible chromatin regions in NFKB1 mutation carriers compared to WT. Dots represent individuals included in the analysis, red indicates affected NFKB1 mutation carrier, blue unaffected NFKB1 mutation carrier, and black WT. Color on the heatmap indicates region with maximum (red) or minimum (blue) open chromatin. Preferentially open regions amounted to 168 in NFKB1 mutation carriers and 381 in WT. **b)** Cis-regulatory sequences associated with transcription factor binding motif (TFBM) regions of the NFKB1mutation carrier cluster or WT cluster. Letter size indicates nucleotide frequency. Numbers indicate significance of motif occurrence (binomial test). **c)** Heatmap including hierarchical clustering of 1396 differential accessible chromatin regions in affected NFKB1 mutation carriers compared to WT. Red dots indicate affected mutation carriers, and black dots WT. Preferentially open regions amounted to 479 in affected NFKB1 mutation carriers and 917 in WT. **d)** Cis-regulatory sequences associated with TFBM regions of the affected NFKB1 mutation carrier cluster or WT cluster. Letter size indicates nucleotide frequency. Numbers on the right indicate significance of motif occurrence (binomial test). **e)** Clustered heatmap for the 377 differentially accessible chromatin regions in unaffected NFKB1 mutation carriers compared to WT. Blue dots indicate unaffected NFKB1 mutation carrier and black dots WT. Blue color on the heatmap represents preferentially open regions, and red preferentially closed regions. Preferentially open regions amounted to 248 in unaffected NFKB1 mutation carriers, and 129 in WT. **f)** Cis-regulatory sequences associated with TFBM regions of the unaffected NFKB1 mutation carrier cluster or WT cluster. Letter size indicates nucleotide frequency. Numbers indicate significance of motif occurrence (binomial test). **g)** Heatmap including hierarchical clustering of 392 differential accessible chromatin regions in affected compared to unaffected NFKB1 mutation carriers. Red dots represent affected mutation carriers. Blue dots represent unaffected mutation carriers. Preferentially open regions amounted to 150 in affected and 242 in unaffected NFKB1 mutation carriers. **h)** Cis-regulatory sequences associated with TFMB regions of affected NFKB1 mutation carrier cluster or unaffected NFKB1 mutation carrier cluster. Sequence logos and P-values reflect the significance of motif occurrence are shown next to the corresponding motif.

Super-enhancers (SE) are regulatory elements characterized by high levels of chromatin modification^59^. These non-coding elements distribute regulatory information within the 3D nuclear chromatin architecture creating SE-promoter contacts that modulate a cell-specific gene expression^60^. NF-κB has a role in the activation of SE^61,62^. To study the super-enhancer landscape of NF-κB target genes, we looked for chromatin accessible regions on *NFKB1* mutation carriers compared to WT (Fig 3). The majority of preferentially accessible regions were on the unaffected group (n=15), followed by WT (n=10) and affected (n=4). In the unaffected group, key regions for B cell proliferation and differentiation were preferentially open (*BCL2, BCL3, CD83, FOSB, NR4A2* and *IRF1*), but we also found open regions for NF-κB negative regulators, *TRAF1* and *TNIP1*. In the WT group, we identified *BCL3, FOSB, IRF1*, and *NR4A2* being preferentially open. These factors act synergistically with NF-κB to activate transcription upon BCR stimulation. The negative regulator *NFKBIA* was also open in this group. Finally, in the affected group, *FOSB, IRF1*, and *NR4A2* were preferentially open, as well as the negative regulator *NFKBIA*. Previous studies have associated SE with NF-κB target genes *in vitro* and in mouse models^61,63,64^. Our analysis indicates that in NF-κB mutated cells chromatin opening at SE is disturbed. This might be an additional factor influencing the TF-DNA binding that causes variation in NF-κB target gene expression.

**Fig 3.**
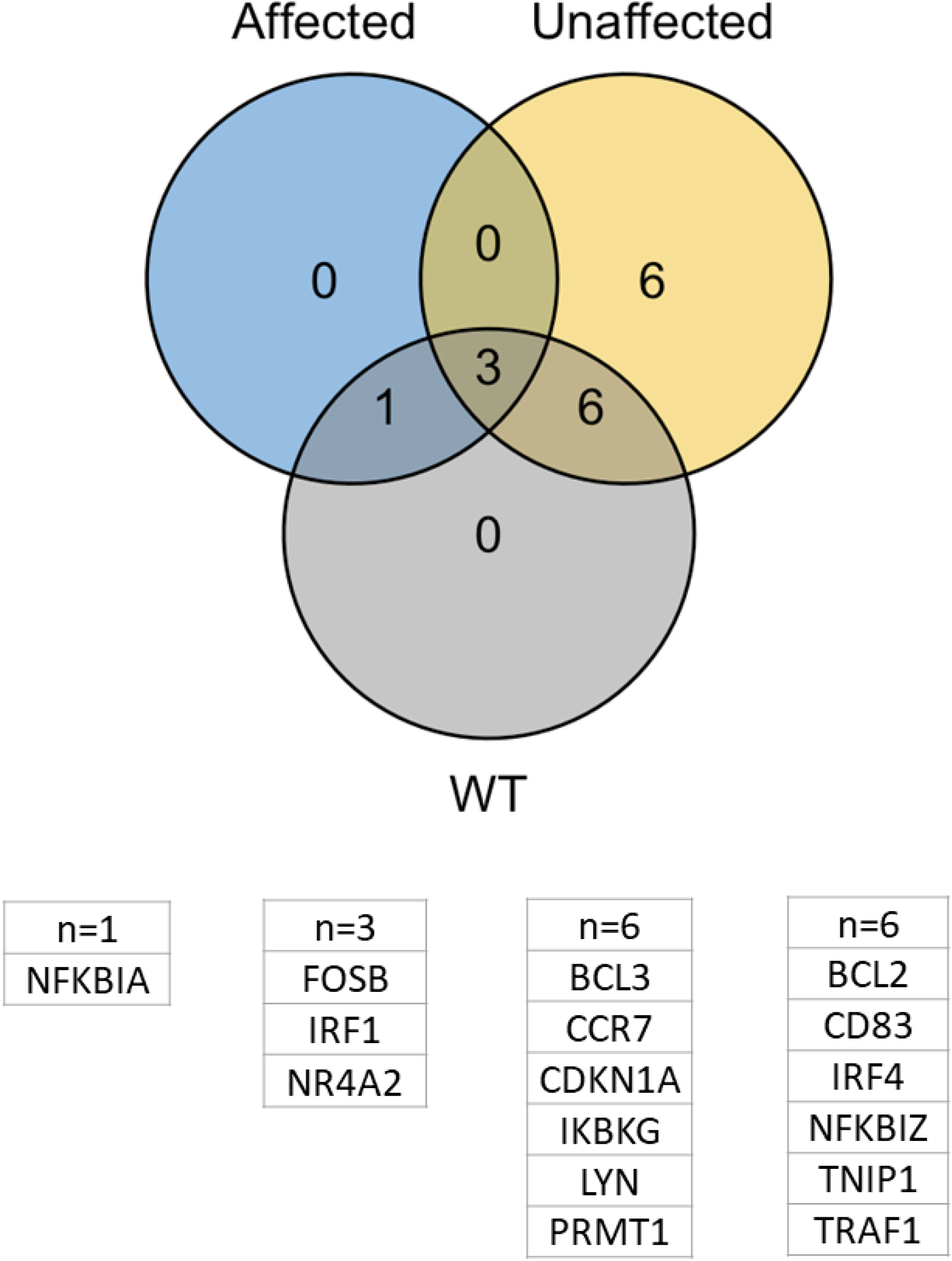
Chromatin accessibility at superenhancers is disturbed on stimulated naïve B cells from NFKB1 patients. Venn diagram representing differentially accessible regions of NFKB target genes at super-enhancer regions. Differentially accessible regions from NFKB1 mutation carriers (affected and unaffected) and WT are represented. Associated gene names are listed in columns below the diagram

Taken together, our data suggest that a network of unique chromatin accessibility profiles result in differences on NFKB1 mutated cells, affecting B cell function and differentiation upon activation.

### Aberrant expression of NF-κB target genes is independent from changes of the methylation pattern in four distinct B cell subsets from *NFKB1* patients

To further explore underlying mechanisms of *NFKB1*-disease in our cohort, we performed in parallel DNA methylation analysis by RRBS and transcriptional profiling by bulk RNA-seq in four distinct unstimulated B cell subsets (unstimulated naïve-, IgM memory-, switched memory-, and CD21low-B cells) derived from three *NFKB1* mutation carriers (FR95, FR103 and FR302), and four *NFKB1* wild-type individuals (C1, C2, C3 and C4). All mutation carriers harboured a pathogenic variant leading to haploinsufficiency (Tables 1 and 2).

We performed transcriptome analysis with a cut-off p-value of <0.05, and an absolute log2-fold change of >0.5. Comparison of WT to *NFKB1* mutation carriers revealed significant expression changes in the unstimulated B cell subsets. In naïve B cells, 1193 genes were differentially expressed (727 downregulated and 466 upregulated), while 33 genes were downregulated in IgM memory-B cells. In addition, 1230 differentially expressed genes were found in switched memory-B cells (869 downregulated and 361 upregulated), and 227 differentially expressed genes in CD21 low-B cells (192 downregulated and 35 upregulated). In *NFKB1* mutation carriers, hallmark gene pathway analysis revealed a predominantly negative enrichment of TGF-beta signaling in all but the CD21low B cell compartment. In naïve B cells, the mTOR signaling pathway, often activated in tumors^65^, was positively enriched, while inflammatory response, hypoxia, and the UV response were negatively enriched (Fig 4a). The role of NF-κB in these processes has been associated with the disruption of cell homeostasis in the disease context^66,67^. In IgM memory- and switched memory-B cells, we found the interferon gamma (IFN-γ) response negatively enriched (Fig 4b and c). Zumakero et al.^68^ demonstrated that IFN-γ induces an epigenetic reprograming of B cells, which results in increased chromatin accessibility of transcription binding motifs for NF-κB, IRF4 and BLIMP1 and thus promotes the formation of antibody secreting cells. However, in CD21low B cells we found merely positive enrichment (Fig 4d). These data are consistent with enriched CD21low populations in patients with chronic immune activation^17^. Moreover, KEGG pathway analysis showed increased *BTK* expression in unstimulated naïve B cells and switched memory B cells. In depth analysis can be found in the supplementary material Figures S7-S12. Our findings highlight, how *NFKB1* mutations profoundly alter B cell biology through the activation or repression of NF-κB target genes.

**Fig 4.**
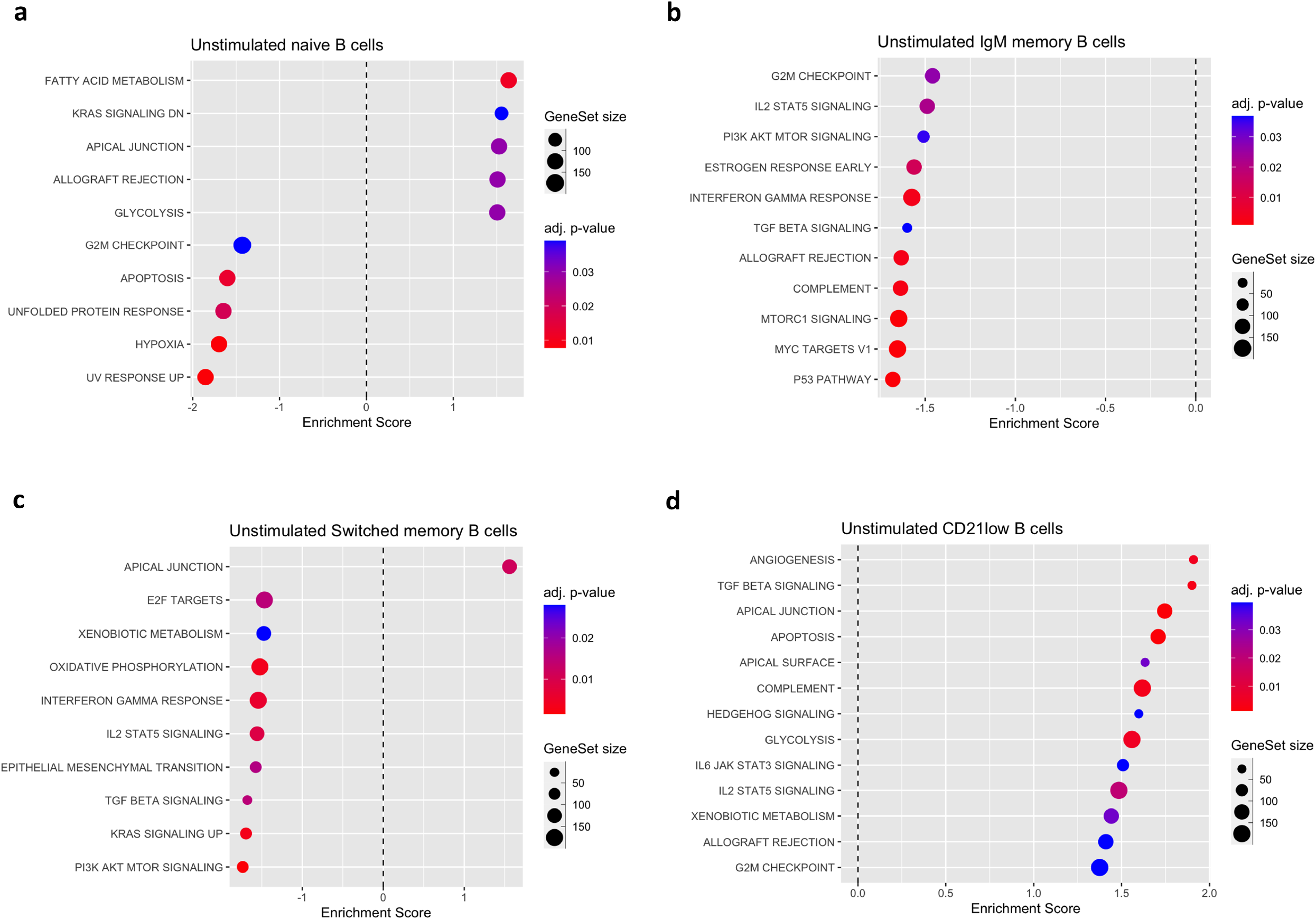
Gene set enrichment analyses show perturbations within the NFKB signaling network on unstimulated B cells across different subtypes. **a)** Dot plot pathway enrichment map showing significant up- and down-regulated genes on unstimulated naïve B cells, **b)** Negative enrichment of pathways on IgM memory B cells when compared to WT **c)** Up- and downregulated gene pathways on switched memory B cells when compared to WT, **d)** Positive enrichment of pathways in CD21-low B cells. The GSEA analysis was performed using the fgsea package in R, where adjusted p-value <0.05 was considered significant. All differentially expressed genes were used for the analysis. Graphs include the gene count (number if DEG in pathway), and the adjusted p-value.

Subsequently, we characterized the methylation profiles of all four unstimulated B cell subsets. Differences in methylation patterns between *NFKB1* mutation carriers and WT were considered relevant when loci (β-values) were 10% or more differentially methylated, with an FDR of <0.05. We observed gradual global demethylation on B cells from the naïve to the memory compartment in healthy individuals, as previously reported^69–72^. Differences in the global methylation pattern were evident at the transition from the naïve, to the IgM-, and switched memory compartments. Patients showed higher methylation levels when compared to WT (Fig 5a). In healthy controls, 14,349 DMRs were hypomethylated in the transition from naïve to IgM memory B cells and 5,028 DMRs in the transition from IgM to switched memory compartment. In contrast, *NFKB1* mutation carriers showed impaired demethylation. We observed 9,137 hypomethylated DMRs at the naïve to IgM memory stage, and 1,785 at the IgM memory to switched memory compartment (Supp.TableS4). These findings corroborate the important role of methylation during B cell differentiation and suggest a contribution of aberrant methylation events to B cell dysfunction.

**Fig 5.**
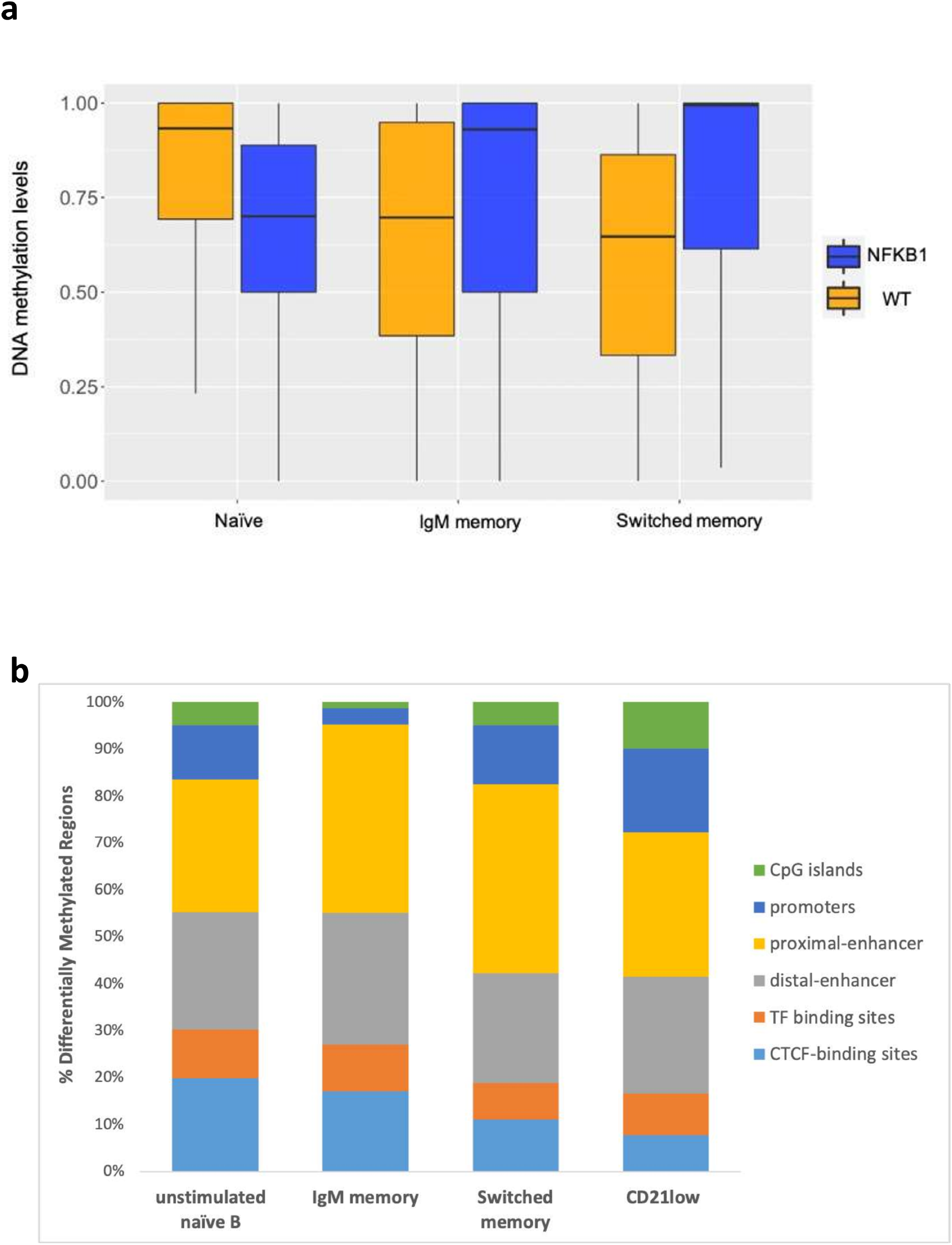
DNA methylation is impaired in the transition from naïve to memory compartments in unstimulated B cells from NFKB1 patients. **a)** Box plot depicting DNA methylation levels of DMRs on the different B cell subsets of NFKB1 mutation carriers(n=3) and WT (n=4). The lower and upper limits of the colored boxes represent the first(Q1) and third quartiles(Q3), respectively. The black horizontal line represents the media. Whiskers indicate the minimum and maximum values of the data set, and outliers are plotted as gray points. To test for significant differences Two-sided Wilcoxon test was used (* p-value ≤ 0.05, NS: not significant). **b)** Location proportion of DMRs across four different cell subtypes in the context of CpG islands, promoters and regulatory regions.

We then focused on differential methylation distribution across various regions (Figure 5b). The majority of DMRs were located at gene regulatory regions, particularly at enhancers, CTCF-, and transcription factor binding sites. Enhancers are known for dynamic methylation patterns^73,74^. During B cell development, around 40% of genes with methylated enhancers are included in the regulatory process of gene expression ^71,75,76^. Several papers suggest that DNA-methylation affects transcription factor binding, indicating that a removal of methylation allows transcription factors to bind within the landscape of hypomethylated regulatory regions^77–80^. Particularly, CTCF has been reported to have a highly specific response to low levels of methylation^81^.

At CpG islands, we identified 470 DMRs (151 hypomethylated and 319 hypermethylated) in unstimulated naïve B cells, 346 DMRs (142 hypomethylated and 214 hypermethylated) in IgM memory B cells, 925 DMRs (346 hypomethylated and 579 hypermethylated) in switched memory B cells, and 963 DMRs (299 hypomethylated and 664 hypermethylated) in CD21-low B cells. When looking specifically at promoter regions, 78 DMRs were identified in unstimulated naïve B cells (43 hypomethylated and 35 hypermethylated), 58 DMRs in IgM memory B cells (25 hypomethylated and 33 hypermethylated), 170 DMRs in switched memory B cells (89 hypomethylated and 81 hypermethylated), and 178 DMRs in CD21-low B cells (82 hypomethylated and 96 hypermethylated). However, none of these DMRs were linked to NF-κB target genes, which was unexpected, since DNA methylation at promoter regions is generally linked to the maintenance of gene silencing^82^ and various NF-κB target genes were found with reduced transcriptional activity.

During B cell differentiation and upon antigen encounter, peripheral mature B cells undergo somatic hypermutation (SHM) and class switch recombination (CSR), which are both associated with a high cellular replication rate and extensive demethylation^83,84^. NF-κB contributes to CSR by binding to the 3’ enhancer, a locus control region located downstream of *IGHA1* ^85,86^. However, methylation of *IGHA1* remained unchanged in *NFKB1* patients. Instead, we found hypermethylation at promoter regions of *IGHD* in unstimulated naïve B cells and of *IGHV4-28* and *IGLV9*-49 loci in IgM memory B cells (Table 4).

**Table 4.**
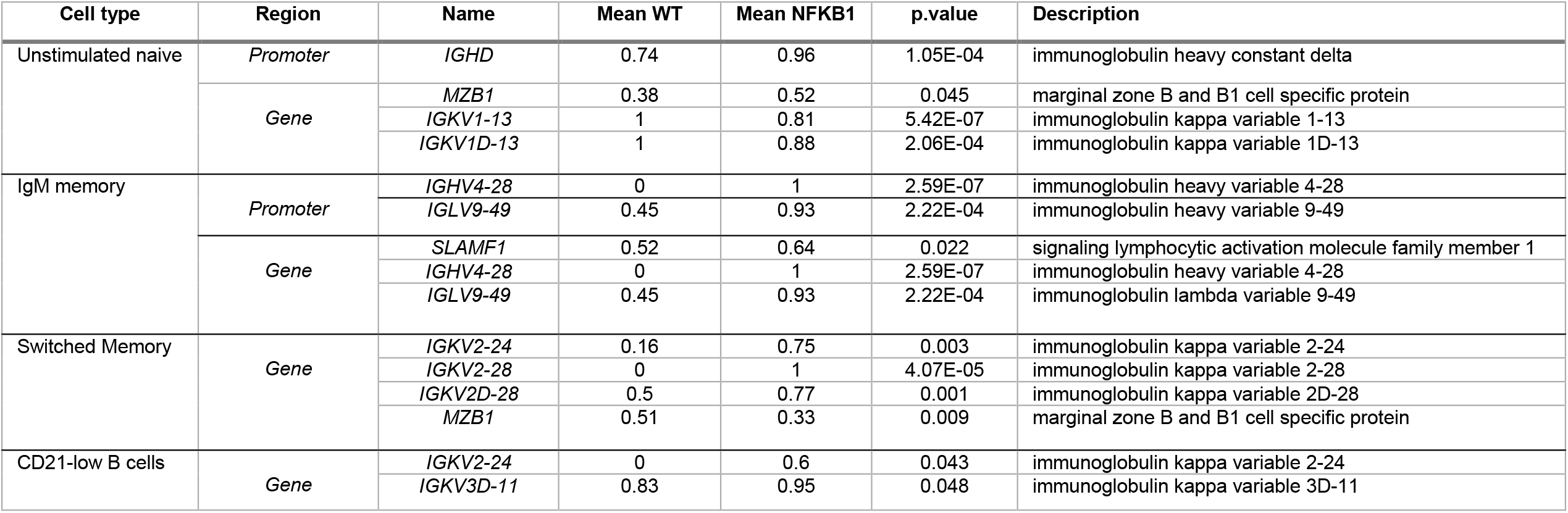
Selected DMR among different B cell subtypes

Plasma cell differentiation is a coordinated process that involves changes in gene expression, and hence gene regulation^87^. When looking at the nearest gene to the DMRs, we found several genes with a role in B cell development dysregulated. Among others, *NFKB1* patients compared to WT showed higher methylation levels of *MZB1* in naïve B cells and lower levels in the switched memory compartment. *MZB1* is expressed in marginal zone B cells and it is highly upregulated during plasma cell differentiation^88,89^. In addition, we found higher levels of methylation of *SLAMF1* in *NFKB1* patients in comparison with healthy individuals in IgM memory B cells. *SLAMF1* is part of the signaling lymphocytic activation molecule family and has a binding site for *NF*κ*B*. In B cells it is upregulated following activation, and plays a role in T and B cell co-stimulation^90^. Increased expression has been reported in patients with autoimmune thyroiditis, lymphoid leukemias and lymphomas ^91–93^.Moreover, we found DMRs at the immunoglobulin chain locus of *NFKB1* patients when compared to WT (Table 4). Expression and rearrangement of immunoglobulin genes takes place during B cell differentiation^94^. The contribution of aberrant DNA methylation to antibody production has been suggested by increased methylation levels in patients with CVID and phenotypically discordant monozygotic twin studies ^72,95^. Accordingly, our data supports the notion that dysregulated DNA methylation also contributes to B cell dysfunction in patients with *NFKB1* mutations. In line with the above findings, the limited expression of negative regulators of NF-κB signaling at the transcriptome level may be a key constituent for the observed hyperinflammatory immune dysregulation in *NFKB1*-mutated patients. Additionally, p105/p50 insufficient patients display altered methylation patterns in distinct B cell subsets, which might contribute to the defective B cell function.

### Proteome analysis of stimulated naïve B cells from *NFKB1* mutation carriers also shows dysregulation in the B cell function

To elucidate effects of *NFKB1* mutations on the protein level, we examined the proteome of CD40L and IL21-stimulated naïve B cells from three affected and three unaffected *NFKB1* mutation carriers and five healthy donors. All individuals had also been included in the transcriptome analysis. 4,666 differentially expressed proteins were detected in the *NFKB1* mutation carriers with cutoffs at >0.5 absolute log2 fold change and <0.05 adjusted p-value (Supp. figure S13).In contrast to the transcriptome data, pathway analysis using Hallmark genes revealed IFN-α and -γ as most enriched in affected *NFKB1* mutation carriers. IFN-α lowers the threshold for B cell antibody responses^96^, while IFN-γ inhibits B cell proliferation during maturation^97^. The pathways for epithelial-mesenchymal transition, coagulation, and UV-response were reduced (Figure 6a), all of which are pathways known to be activated by *NFKB1* ^98,99^. The same pathways were observed to be decreased in unaffected *NFKB1* mutation carriers. However, in unaffected patients PI3K-Akt-mTOR signaling was most enriched (Figure 6b). The PI3K-Akt-mTOR axis is essential for normal B cell metabolism, proliferation and development ^100^.

**Fig. 6.**
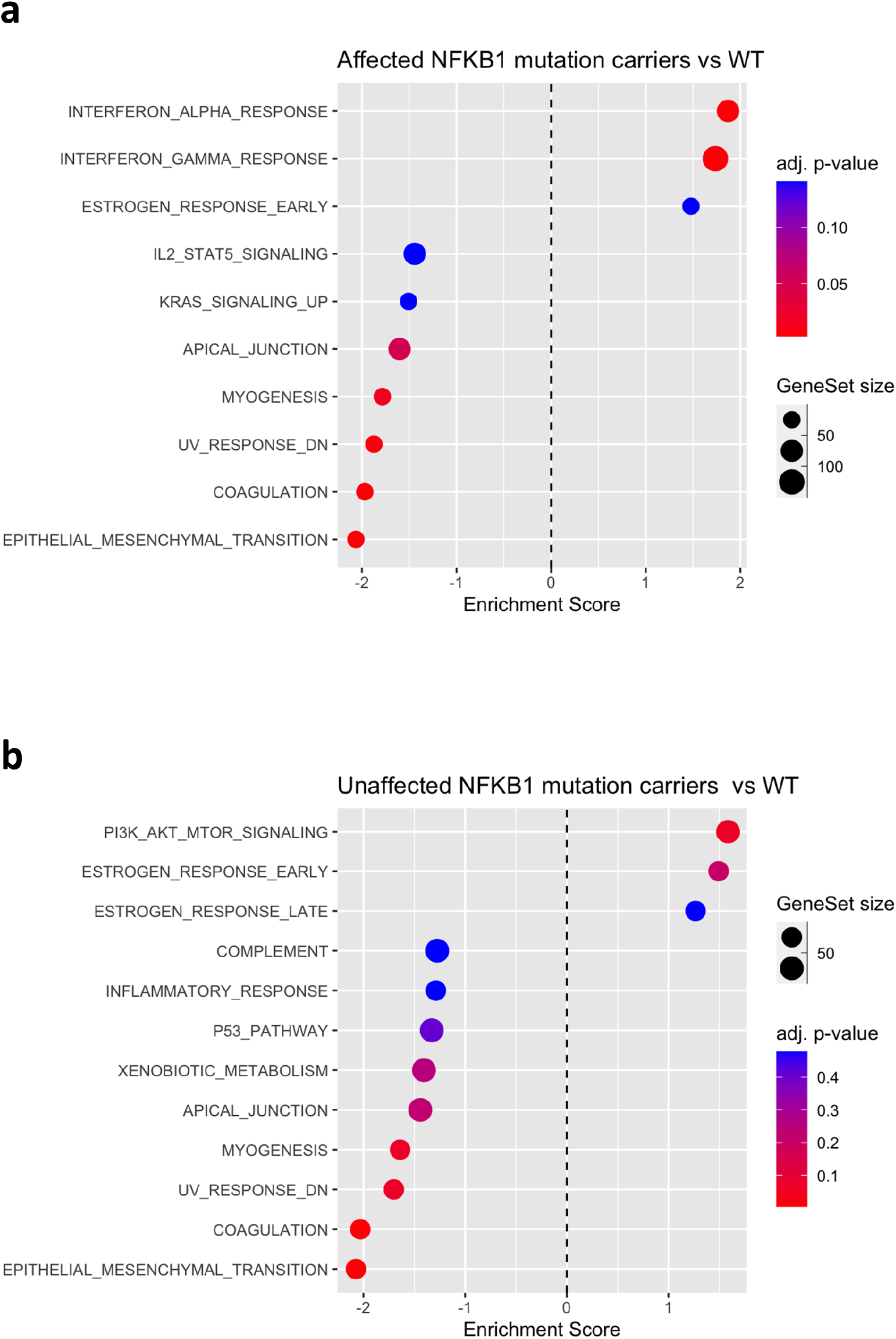
The proteome of stimulated NFKB1 insufficient cells also reflects B cell dysfunction. Dot plots representing hallmark pathways. **a)** Hallmark pathways associated with increased (right) or decreased (left) differentially expressed proteins on affected mutation carriers when compared to WT. All categories are statically significantly enriched (adjusted p-value <0.5) **b)** Unaffected mutation carriers statistically significant enriched Hallmark pathways (adjusted p-value <0.5) associated with increased (right) or decreased (left) differentially expressed proteins, when compared to WT, are depicted in the dot-plot. Graphs include the gene count (number of DEP in pathway), and the adjusted p-value.

When correlating log2-fold changes for differentially expressed genes with their corresponding differentially expressed proteins, we found a statically significant positive correlation (p=0.07, r=0.06) for affected and unaffected *NFKB1* mutation carriers compared to WT (Figure 7a). However, when we separating the groups according to disease status (affected or unaffected) and comparing them to WT, no correlation was observed (Figures 7b and 7c). A weak correlation between mRNA transcripts and proteins could be explained by translation regulation, and post-transcriptional protein modifications and degradation^101^, which became evident only after we analyzed the groups according to disease status.

**Fig 7.**
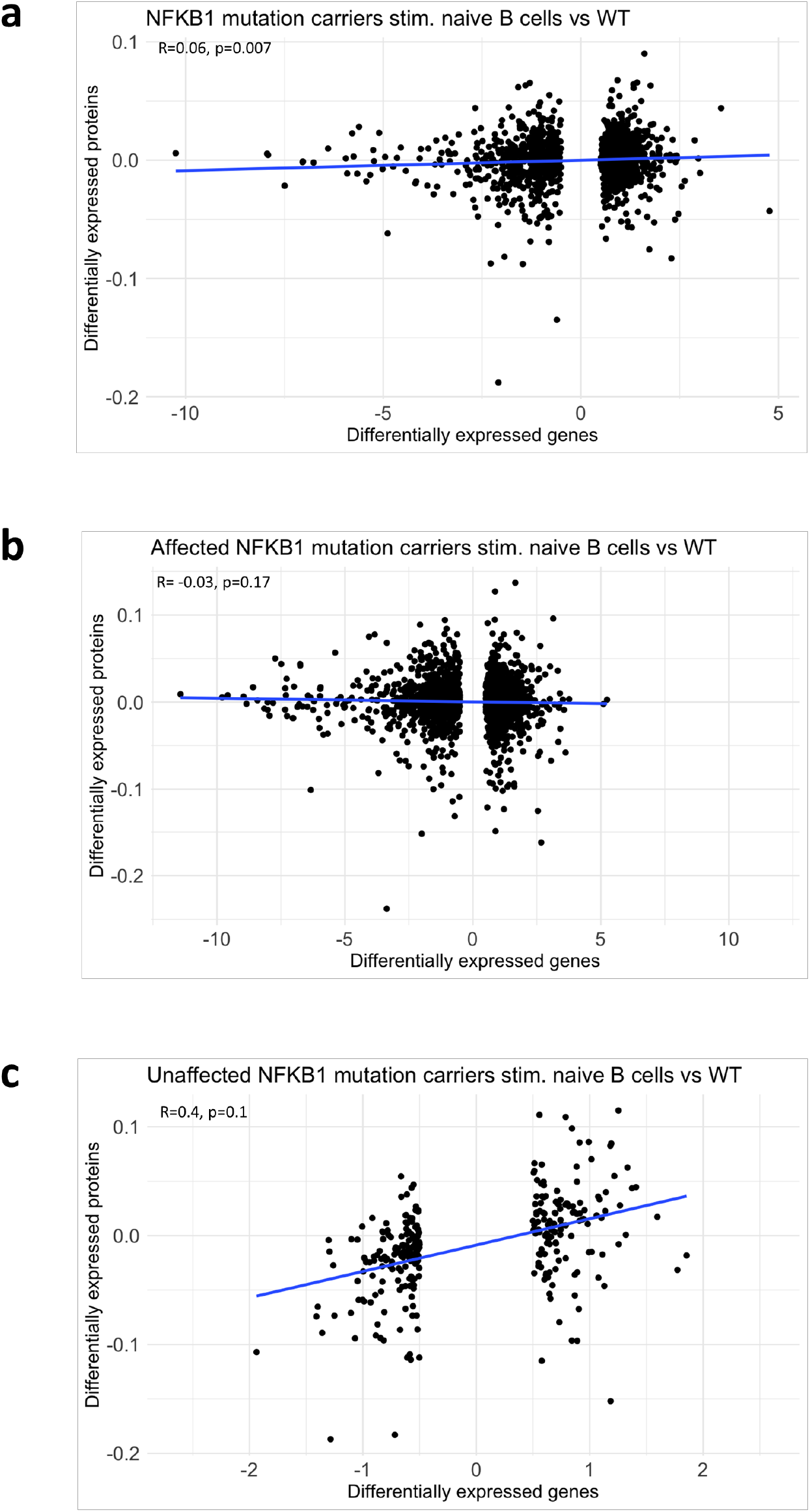
Low correlation between differentially expressed genes and proteins on stimulated naïve B cells in NFKB1 mutation carriers. Scatter plots showing the correlation between **a)** NFKB1 mutation carriers compared to WT, **b)** Affected NFKB1 mutation carriers compared to WT, and **c)** Unaffected NFKB1 mutation carriers compared to WT. Scatterplot depicts the Pearson correlation coefficient (r) of differentially expressed genes (log2 fold change ≥0.5), and differentially expressed proteins (alpha = 0.05 and an absolute log2 fold change of 1.5) from the comparison between NFKB1 mutation carriers and WT.

### Multi-omics analysis of *NFKB1*-insufficient B cells reveals accessible chromatin regions with incomplete demethylation

Given the essential role of NF-κB signaling in various physiological processes and the severe consequences of potential dysregulation, NF-κB activity requires tightly adjustable control mechanisms. Since active transcription factors bind to chromatin in order to modulate target gene expression^102^, chromatin accessibility and epigenetic modifications, such as DNA methylation, can determine the duration and strength of NF-κB binding and therefore the transcriptional output^103,104^. To explore the relationship between epigenetic changes and differential gene expression in *NFKB1* mutation carriers, we performed an association analysis of DNA methylation levels and preferentially open chromatin regions, with differentially expressed genes and differentially expressed proteins obtained by comparing affected *NFKB1* mutation carriers to WT control individuals (Figures 8a and 8b). While hypomethylation at promoters and preferentially open chromatin regions facilitates transcription, hypermethylation generally associates with lower accessibility. Inspecting our data for such conditions, we found two genes, *BAX* and *SERPINE2*, to have increased expression, open chromatin and hypomethylated promoters. *TRAF3, TRAF6, TP53BP2, STAT5A, RIPK2, NFKBIA, BCL2, ATM, TNFAIP3, NFKBIZ, CD83, and CD40* showed promoter hypomethylation and open chromatin, although transcription was reduced. In contrast, *CDKN1A* had promoter hypermethylation, closed chromatin and reduced expression, while *TNIP1* had promoter hypermethylation, open chromatin and reduced expression. *TRAF1, RELB, TNFAIP2, NFKBIE* and *REL*, had a reduced transcription, with hypomethylated promoters but closed chromatin. Interestingly, *NFKB1* itself had a preferentially open chromatin and a hypomethylated promoter region, although its transcripts were underrepresented. This observation indicates that the mutant transcripts might undergo accelerated decay, as previously suggested by Fliegauf et al.,^1^ which, in addition to protein loss due to devastating defects, contribute to the p105/p50 haploinsufficient condition in patient-derived cells.

**Fig 8.**
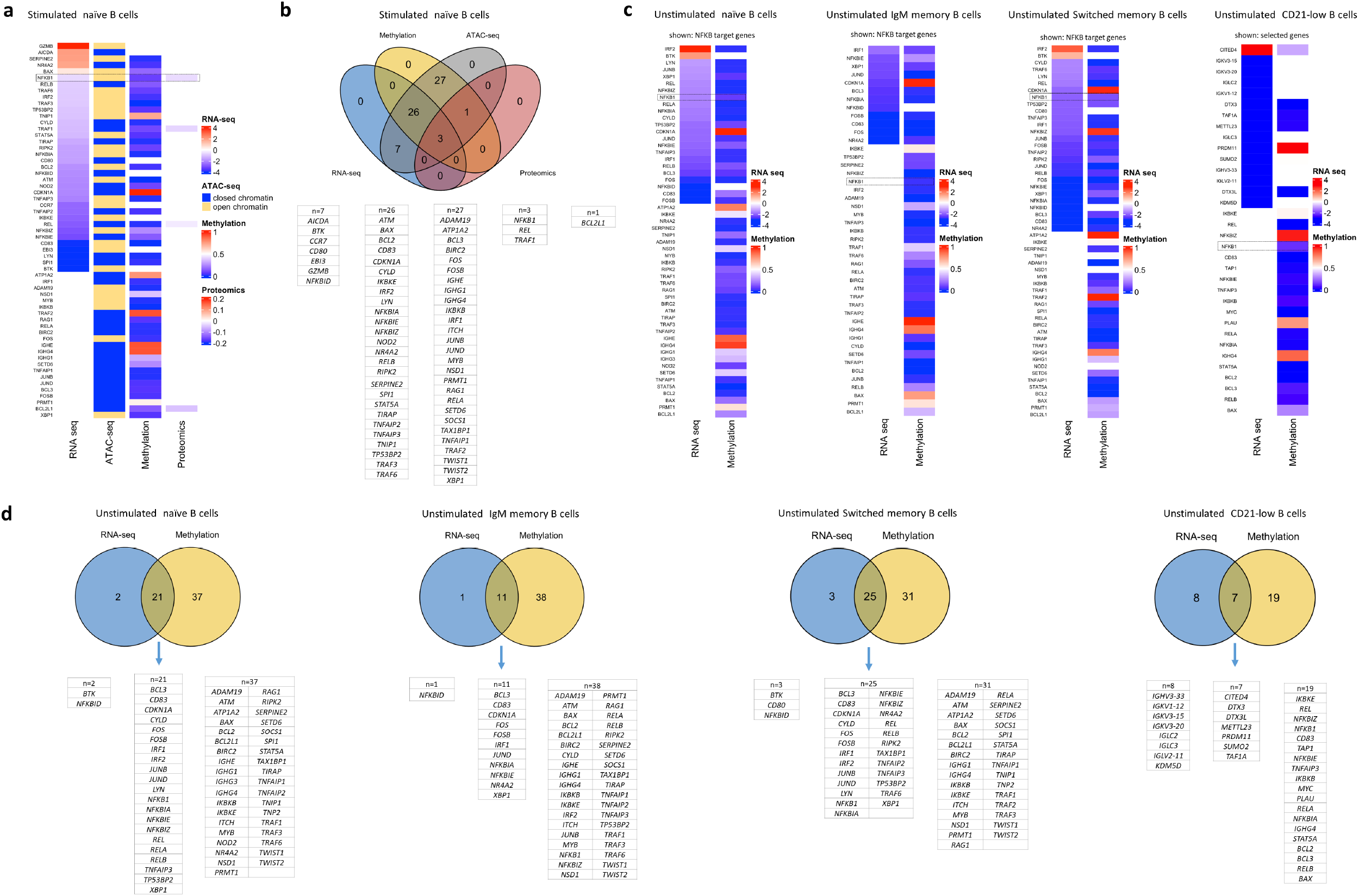
Decrease expression of NFKB target genes in different omic levels on stimulated and unstimulated B cells from NFKB1 patients. **a)** Heatmap representation showing association between transcriptome, chromatin accessibility, methylome, and proteome from affected NFKB1 mutation carriers compared to WT in stimulated naïve B cells. For RNA-seq data, differentially expressed genes obtained from the comparison between NFKB1 mutation carriers to WT are shown. Color indicates log fold change were blue and red represent a decrease, and increased expression, respectively. The cutoff for differential expression was set at p<0.05 adjusted p-value and 0.5 for log2 fold change. For ATAC-seq, preferentially open or closed regions from *NFKB1* mutation carriers are shown. Blue bars represent preferentially closed and yellow bars preferentially open chromatin regions. Methylation data from NFKB1 mutation carriers are depicted. The median methylation is represented as a beta-value from 0 to 1 (0 to 100% methylation) at promoter regions. Color represents methylation values, where red is 100% and blue 0% methylation, respectively. For proteomic data, protein ratios were log-2 transformed. Differentially expressed proteins (alpha = 0.05 and an absolute log2 fold change of 1.5) from the comparison between NFKB1 mutation carriers and WT are represented. Color represents log fold change, where blue and red represent a decreased and increased expression, respectively. **b)** Venn diagram showing a supervised analysis of NFKB target genes in stimulated naïve B cells. Diagram represents the overlap between differentially expressed genes and differentially expressed proteins comparing affected mutation carriers to WT, plus preferentially open chromatin regions and methylation levels in affected NFKB1 mutation carriers. Associated gene names are shown in the columns below the diagram. **c)** Heatmap showing association between transcriptome and methylation levels on unstimulated B cells. A supervised analysis for NFKB target genes for naïve B-, IgM memory, switched memory-, and CD21low-B cells is represented. Additionally, for CD21-lowB cells, selected genes are shown. For differentially expressed genes data were obtained from the comparison of NFKB1 mutation carrier vs WT. Log2-fold changes values are indicated, cutoff for dysregulation was p-value <0.05. Blue and red color indicate decrease or increase expression, respectively. Methylation levels from NFKB1 mutation carriers are depicted. The median methylation is represented as a beta-value from 0 to 1 (0 to 100% methylation) at promoter regions. Blue color indicates 0% methylation, and red 100% methylation. **d)** Venn diagram representing a supervised analysis of NFKB target genes, between methylation levels and differentially expressed genes on unstimulated naïve-, IgM memory- and switched memory B cells. Additionally, for CD21-low B cells, selected genes are shown. Differentially expressed genes were obtained from the comparison between NFKB1 mutation carriers to WT (log2 fold change ≥0.5, and adjusted p-value <0.5). Methylation values from NFKB1 mutation carriers are represented. Associated gene names are listed in columns below the diagram.

To integrate methylation and transcriptome data of unstimulated cells, we analyzed downstream genes of the NF-κB signaling pathway in all four B-cell subtypes (Figures 8c and d). We identified hypomethylation and lower expression levels of members of the IRF (*IRF1*) and AP-1 (JUNB, *JUND, FOS, and FOSB*) families of transcription factors in the naïve-, IgM memory-, and switched memory B cell compartments. Among the IRFs, IRF1 can associate with members of the NF-κB/Rel family, generating complexes that synergistically activate transcription^105^. It has been suggested that NF-κB and Fos/Jun have synergistic mechanisms of gene regulation^106^. *IRF2* was hypomethylated and showed increased expression in naïve B cells and switched memory B cells. IRF2 is an antagonistic transcriptional repressor of IRF1, and interferes with NF-κB activation^107^. Among the genes encoding negative regulators of NF-κB, we found hypomethylation and lower transcript levels for *NFKBIA, NFKBIE, NFKBID* and *TNFAIP3* in naïve B cells, for *NFKBIA* and *NFKBIE* in IgM memory B cells, and for *NFKBIA* and *TNFAIP3* in switched memory B cells. Other members of the NF-κB family (*REL, RELA* and *RELB*) were hypomethylated and their transcripts were reduced, which was observed for *NFKB1* in unstimulated naïve B cells and switched memory B cells. Moreover, *BCL3* was hypomethylated and lower expressed in naïve- and switched memory-B cells, respectively. *BCL3*, a nuclear member of the I-κB family, can enhance transcription of NF-κB target genes by binding to p50 homodimers^108^.

In addition, *CD83*, an activation marker of B cells in the light zone of the GC reaction^109,110^, was hypomethylated and had reduced expression in IgM memory B cells and switched memory B cells. *XBP1*, a transcription factor that enables high immunoglobulin expression and changes associated with terminal B cell differentiation into plasma cells, was hypomethylated and its transcripts were reduced in IgM memory- and switched memory B cells. In the CD21-low compartment we found increased expression and hypomethylation of *CITED4* (CBP/p300-interacting transactivator 4). NF-κB engages CBP/p300 and its histone acetyltransferase activity for transcriptional activation of the IL6 promoter^111^. Moreover, we identified transcripts of genes coding for regulators of the epigenetic machinery, such as methyltransferase *METTL23*, lysine demethylase *KDM5D*, and *PRDM11*, which enables chromatin binding, to be underrepresented in the CD21-low B cell compartment. Additionally, transcripts of *TAF1A* (TATA box binding protein associated factor), genes with a role in ubiquitylation (*DTX3, DTX3L, SUMO2, USP9Y*), and immunoglobulin genes (*IGKV3-15, IGKV3-20, IGKV1-12, IGHV3-33, IGLV2-11*) also showed decreased levels in CD21-low B cells. In-depth analysis of different comparisons can be found in the Supplementary Material Figures S14-17.

Our results demonstrate a compromised B cell function on *NFKB1*-mutated cells in different omics dimensions. First, we found a reduced expression of negative NF-κB pathway regulators. Second, we showed that regions encoding crucial factors for B cell proliferation and differentiation had changes in chromatin accessibility. In addition, our results indicate that *NFKB1*-mutated B cells have aberrant DNA methylation patterns. We also detected changes in protein expression levels linked to B cell differentiation and maturation. Overall, our findings provide evidence for overlapping layers of dysregulation, which collectively dictate aberrant gene activation profiles and changes in the NF-κB network. Following these results, it is likely that direct perturbations within the NF-κB signaling pathway, added to a blurred crosstalk with other families of transcription factors, and changes in epigenetic regulation, explain the aberrant B cell differentiation, hypogammaglobulinemia, and autoimmunity, all observed in our patients. Moreover, the imbalance of the negative feedback loop embedded in the NF-κB system may perpetuate inflammation and autoimmunity.

## Discussion

Pathogenic mutations in *NFKB1* have been associated with various phenotypes ranging from antibody deficiency to immune dysregulation^4,5,7,24^. Given that the NF-κB signaling network plays a central role in many cellular processes, impaired immune functions might originate not only from deleterious expression of the transcription factor, but also from delayed, accelerated or prolonged downstream effects. In this study, we present a multi-omics characterization of *NFKB1* (p105/p50) insufficiency in B cells, involving WES, RNA-sequencing, ATAC-sequencing, methylation, and proteomics. It demonstrates a direct relationship between affected patients and aberrant chromatin accessibility, epigenetic, and transcriptional alterations in NF-κB target genes, including members of the network of negative feedback loop that ensure the termination of the NF-κB response.

Based on WES data, we identified *NFKB1* mutation carriers harboring other variants with low allele frequency in NF-κB target genes and genes causing inborn errors of immunity. Due to the complexity of the NF-κB signaling pathway, we underscore the possibility that these variants additionally modify NF-κB signaling at various stages; hence, they might act synergistically or antagonistically, and therefore influence disease presentation and severity. This could potentially contribute to the very variable disease penetrance and expressivity observed in patients with *NFKB1* insufficiency. That this concept is a viable hypothesis has been demonstrated in selected publications such as in Sic et al.^112^.

We showed that *NFKB1*-insufficient B cells display widespread transcriptome changes that are present in stimulated and non-stimulated naïve B cells, IgM memory B cells, switched memory B cells, and to a lower extent in CD21low B cells. The activation of NF-κB is tightly regulated and requires a cascade of post-translational modifications through phosphorylation and ubiquitination events that culminate in the activation of the signalling pathway^104^. Interestingly, the ubiquitination regulators TRAF2, TRAF6, cIAP1/2, and TAB proteins had reduced expression levels in mutation carriers. Moreover, both TNF-α and IL-1 proinflammatory responses are mediated by NF-κB signaling^18^. We found a repression of genes related to TNF-α signaling in *NFKB1* mutation carriers, however, we cannot say whether this is directly caused by the mutation in *NFKB1* or whether this is a regulatory effect by the B cells in order to limit NF-κB-mediated hyperinflammation.

The NF-κB signalling pathway contains homeostatic feedback mechanisms that limit and restore its inducible activation^55^. In mutation carriers, we found transcripts of genes encoding for negative regulators to be underrepresented^18^: *TNFAIP3* codes for the deubiquitinase A20 which removes polyubiquitin chains from TNF receptor-associated factor 6 (*TRAF6*); and *TRAF1* which interferes with the linear ubiquitination of NEMO. Also, *TNIP1*, which encodes a component of the A20-ubiquitin editing complex, and *CYLD* which deubiquitinates *TAK1* were found to be downregulated. *CYLD* is a regulatory enzyme with deubiquitylating activity. *CYLD* it is not a direct NF-κB target gene, but controls de deubiquitinylation of NF-κB target genes, including *TRAF2, TRAF6, TAK1, BCL3, IKKY, RIP1, RIGI*, and *TBK1*. Mutations in *CYLD* can be found in patients with familial cylindromatosis, hair-follicle keratinocytes, and nasopharyngeal tumors, suggesting its role as a tumor suppressor^113,114^. In keratinocytes, Massoumi et al.^115^ demonstrated that CYLD inhibits cell proliferation and tumor growth through the block of BCL3-NF-κB dependent signaling. Non-Hodgkin B cell lymphoma and solid organ cancer have been reported in around 16% of *NFKB1* insufficient patients^7^. A reduced expression of *CYLD* could be a clue for the pathogenic mechanisms of malignancy in these individuals, but requires further investigation.

Aberrant inactivation of NF-κB activity is a common factor of autoimmunity, autoinflammation and cancer^116–118^. Germline frameshift mutations in *TNFAIP3*, resulting in A20 haploinsufficiency have also been identified in patients with an autoimmune lymphoproliferative syndrome-like phenotype (ALPS) and T cells from these patients exhibit increased NF-κB activation^119^. Germline loss-of-function in *TNFAIP3* leading to A20 haploinsufficiency was shown to cause Behcet-like autoimmunity^120^, a phenotype which has also been reported in *NFKB1*-mutated individuals. Actually, all affected mutation carriers from our cohort presented with autoimmunity and autoinflammation. Dysregulation of NF-κB downstream inhibition might prevent appropriate attenuation of proinflammatory signaling and could lead to increased inflammation in *NFKB1* haploinsufficiency. This observation is important for the development of potential new drug targets. The ubiquitin-proteasome system plays an important role in the activation of NF-κB. The use of proteasome inhibitors like Bortezomib was shown to be effective in autoimmune disorders such as systemic lupus and myasthenia gravis^121^. Nevertheless, positive feedback mechanisms within the signaling pathway might also contribute to human disease. As TNF-α and IL1 initiate the canonical pathway and also are an NF-κB target genes, it might create a positive feedback loop until signals perpetuating this cascade are eliminated. Thus, targeting TNF-α or IL1 signaling also constitutes a promising therapeutic approach especially in patients presenting with an inflammatory phenotype (anti-IL1) and gastrointestinal involvement (anti-TNF-α) ^122^.

Another group of anti-inflammatory proteins are the suppressors of cytokine signaling (SOCS), which interfere with proinflammatory signaling pathways, such as the JAK-STAT and NF-κB signaling network ^123^. *SOCS1* codifies for a protein that functions as a ubiquitin ligase, leading to polyubiquitination and proteasomal degradation of p65 in the nucleus^124,125^. When we compared affected to unaffected *NFKB1* mutation carriers, we found *SOCS1* to be lower expressed in affected patients. This could add to the disease status of our patients, as less *SOCS1* means more p65 and hence more proinflammatory gene transcription. Especially in concert with the observed increased presence of interferons described above, seen in the proteome of our *NFKB1*-mutated individuals, inhibition of the JAK-STAT signaling pathway may also be beneficial for these patients.

The production of granzyme B in NF-κB1-deficient B cells seems also noteworthy to us: granzyme B is being produced by activated B cells^50^ e.g. when detecting viral antigens^52^. Hence, its production in our patient’s B cells may either point towards undiagnosed chronic viral infection(s) (in this case it would be secondary to the genetic defect), or the *NFKB1*-mutation itself leads to an activated state of B cells (the mutation would be the primary cause for granzyme B expression). This observation also raises the hypothesis that in CVID patients without underlying genetic defect, possibly chronic viral infections may be the underlying culprit?

The expression of *SERPINE2* in CD21-low B cells could indicate that these cells may be the consequence of an insufficient germinal center reaction not only in NF-κB1-insufficient patients, but possibly also in other patients with increased levels of CD21-low B cells, such as in patients with systemic lupus or HIV infection ^126,127^. *SERPINE2* is known to be expressed in cells undergoing homologous recombination DNA repair ^128^. The hypothesis is that during a defective or incomplete GC reaction, in which double strand breaks occur at the human IgG locus on chromosome 14 for class switch recombination, DNA repair seems to be insufficient, giving rise to a constant pool of *SERPINE2*-expressing CD21-low B cells.

At the chromatin level, we detected an enrichment of binding motifs of transcription factors associated with B-cell function in stimulated naïve B cells. For example, CTCF is required to sustain the germinal center transcriptional program to avoid premature plasma cell differentiation^129^. Moreover, BATF and Fra1, a member of the AP-1 family of transcription factors, were enriched. Fra1 is involved in the regulation of follicular B-cell differentiation into plasma cells^130^. BATF controls activation-induced cytidine deaminase (AID) and germline transcripts of the intervening heavy-chain region and constant heavychain region^131^. These findings are in agreement with a recent report on CVID and epigenetic dysregulation in B cells^72^. Epigenetic alterations affecting genes and regulatory elements in memory B cells impair plasma cell differentiation^87,132^ While we did not find DNA methylation changes in naïve B cells, we found demethylation defects occurring from the transition from naïve, to IgM memory-, and to switch memory-B cells. These data are consistent with the observed hypermethylation in gene regions at immunoglobulin loci in IgM memory, switched memory- and CD21low-B cells in our datasets. This is an important observation, because DNA methylation changes are required for somatic hypermutation and class switch recombination^84,133^.

Our omics approach confirmed that chromatin accessibility is a key regulatory mechanism to facilitate transcription factor binding^103^. NF-κB is able to influence the chromatin state through a variety of mechanisms, including the recruitment of components of the general transcription machinery. Frequently, the selective interaction with these co-activators is dependent on specific post-translational modification of NF-κB subunits^134^. We observed that methylation defects may compromise the functionality of the transcriptional machinery, and that aberrant regulation at different layers might be the cause of hypogammaglobulinemia and autoimmunity in these patients. More detailed proteomic data may provide deeper insights into the downstream signaling consequences of *NFKB1* insufficiency. As shown in previous reports^135,136^ the correlation between RNA and protein analysis was low also in our study. NF-κB controls numerous signaling cascades in the immune system. Therefore, it is conceivable that haploinsufficiency mutations affecting also the function of negative regulators of the NF-κB pathway, are leading to autoimmune or inflammatory conditions.

One limitation of this study is that we did not perform experiments in cell lines or in a murine system, and these experiments need to be conducted to move from the descriptive state of associations, as presented here, to the causality and proven NF-κB network dependencies and interactions. Moreover, during the course of this project we learned that single cell analysis, although more expensive, would most likely have been superior to our approach of bulk sequencing of sorted B lymphocyte subsets.

Here, we described the implications of pathogenic *NFKB1*-mutations on B cells on different levels. Our study highlights the additional insights that can be gained by integrating omics data sources. Our analysis provides evidence of hyperinflammation in *NFKB1*-mutated cells caused by the reduced expression of negative regulators. This may open new pathways for the development of personalized targeted therapy. We consider that the observations described here, and the data underlying them, will be a key resource for future in-depth studies of the immune dysregulation associated with defects of the NF-κB signaling network.

## Supporting information

Supp.TableS1_Primer list

Supp.TableS2_ Genetic variants with low AF on NFKB1 mutation carriers

Supp.TableS3_Available dataset per sample

## Abbreviations

ATAC-seq: assay for Transposase-Accessible Chromatin using sequencing
CSR: class switch recombination
CVID: Common Variable Immunodeficiency
FACS: fluorescence activated cell sorting
NFKB1: Nuclear Factor kappa B subunit 1
PBMCs: Peripheral Blood Mononuclear Cells
RRBS: Reduced represented bisulfite sequencing
SE: Superenhancers
TFBM: transcription factor binding motifs
TNF: Tumor Necrosis factor
WES: whole exome sequencing
WT: wild type

## Supplemental information

## Acknowledgements

We thank all affected individuals, their family members and healthy volunteers who participated in this study. We thank Pavla Mrovecova, Katrin Hübscher, Anastasja Maks, and Hanna Haberstroh for excellent technical assistance. We acknowledge the Lighthouse Core Facility for their excellent assistance with cell sorting and flow cytometry. Some samples have been taken from the CCI-biobank, a partner of the Freeze Biobank Freiburg. We thank the Biomedical Sequencing Facility at CeMM for assistance with next-generation Sequencing. The authors acknowledge the support of the Freiburg Galaxy Team: Beatriz Serrano-Solano and Björn Grüning, Bioinformatics, University of Freiburg (Germany) funded by the Collaborative Research Centre 992 Medical Epigenetics (DFG grant SFB 992/1 2012) and the German Federal Ministry of Education and Research BMBF grant 031 A538A de.NBI-RBC.We are grateful to Pascal Schneider for providing CD40L.

## Online resources

Ensembl Genome browser: https://www.ensembl.org/

Boston University Biology: https://www.bu.edu/nf-kb/gene-resources/target-genes/

UCSC Genome browser: https://genome.ucsc.edu/

Galaxy Europe: https://usegalaxy.eu/

GnomAD browser: https://gnomad.broadinstitute.org/downloads

R studio: https://www.rstudio.com/

## Data availability

The exome data sets presented in this article are not publicly available because uploading the data is not part of the participants consent according to Art. 7 GDPR. All other omics data sets (RNA-seq, ATAC-seq, RRBS, and proteomics) have been deposited on SRA database (RRBS, RNA-, ATAC-seq) and ProteomeXchange Consortium via the PRIDE^137^ partner repository.

## Ethics statement

The ethics committee Freiburg approved this project with an effective date of 06.06.2019. In July 2019 the consent form under which patient samples had been collected, was approved under ETK-Nr_354/19. The i-PAD ethics vote carries the number EK-Freiburg_76/19 and the NFKB Study protocol has the approval No. 295/13_200149 and 93/18_191111. Written informed consent to participate in this study was provided by the participants themselves.

## Author contributions

NC-O processed and collected samples, performed methylation, RNA-seq and association analysis, collected clinical data, wrote case reports, designed layout, figures and tables, and wrote the manuscript. NR processed and collected samples, analyzed ATAC and proteome data, designed ATAC data figures and wrote the manuscript. SP-C designed RNA-seq script, summarized data, processed and collected samples, wrote the manuscript. AC-O analyzed and interpreted WES data. MF performed functional testing of the variants and revised the work critically for intellectual content. FZ organized and coordinated ATAC and RRBS sample preparation, and performed preliminary quality control. MG processed ATAC sequence in patient samples. VG processed ATAC and RRBS sequencing on samples. MP measured proteome samples and revised analysis. KW cared for patients, provided clinical and immunological information. MP analyzed and interpreted WES, designed sorting strategy, revised the work critically for intellectual content. EB performed conception and designing of the study, revised the work critically for intellectual content. CB performed conception and designing of the study, revised the work critically for intellectual content. GR performed conception and designing of the study, processed proteome sequencing n samples. BG performed conception and designing of the study, provided and cared for study patients and revised the work critically for intellectual content. All authors contributed to manuscript revision, read, and approved the submitted version.

