## Supplementary Figures for "Integrated Multi-omics Analyses of NFKB1 patients B cells points towards an up regulation of NF-κB network inhibitors"

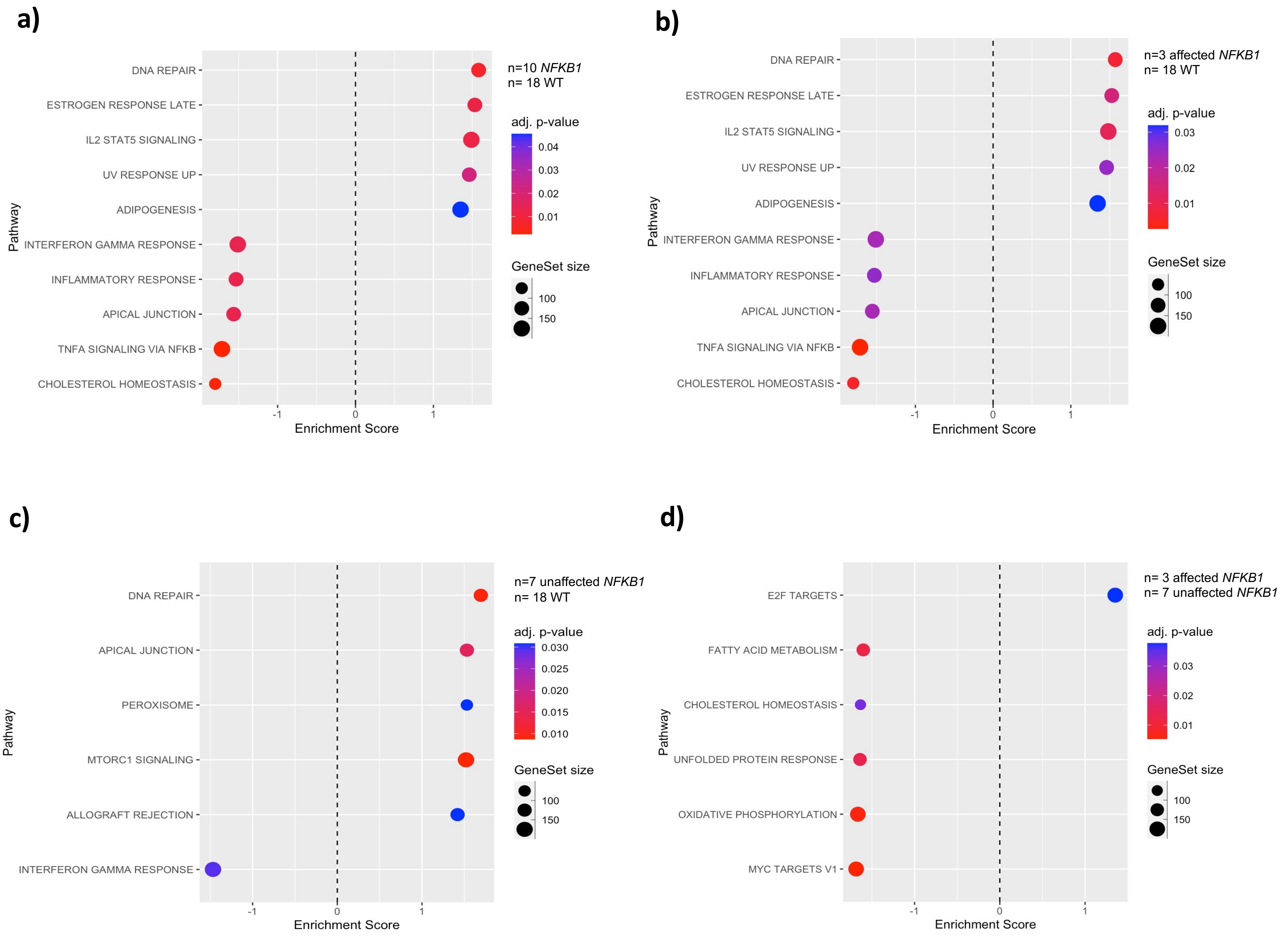


**Supplementary Figure S1**. **Gene set enrichment analyses on stimulated naïve B cells shows downregulation of TNF-α signaling via NF-κB, and Interferon Gamma Response pathways in NFKB1 mutation carriers**. Gene set enrichment analysis representation of differentially expressed genes. All represented categories are statically significantly enriched, with a p value <0.05, as indicated in the color bar located on the right side of each figure. Enrichment score refers to the score between the differentially expressed genes that are associated with a Hallmark pathway term and all genes included in that hallmark pathway. Dot color represents adjusted p values, and dot size indicates the number of times each term is represented in the differentially expressed genes. All differentially expressed genes were used for the analysis. **a)** Enriched pathways in all NFKB1 mutation carriers(3 affected and 7 unaffected) when compared to WT(n=18), **b)** Hallmark pathways up- and down-regulated in affected mutation carriers(n=3) when compared to WT(n=18), **c)** up- and down-regulated gene pathways in unaffected NFKB1 mutation carriers(n=7) when compared to WT(n=18), **d)** Enriched pathways in affected(n=3) when compared to unaffected(n=7) NFKB1 mutation carriers.

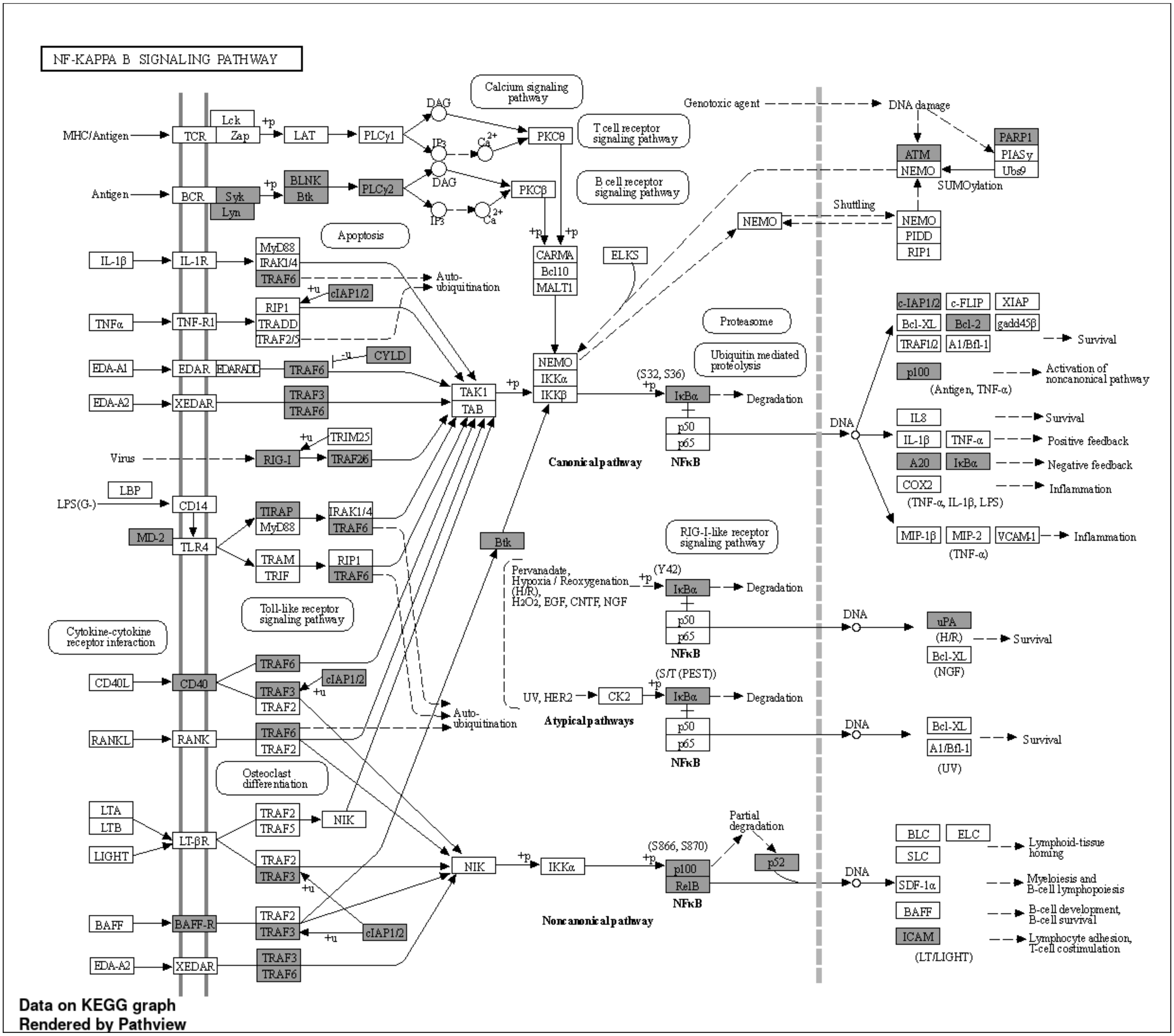


**Supplementary Figure S2**. **Transcriptome analysis revealed decreased expression of genes encoding components of the NF-kB signaling pathway on stimulated naïve B cells from NFKB1 mutation carriers.**
Pathway overview of decreased expression of NFKB target genes comparing NFKB1 mutation carriers (3 affected and 7 unaffected) to WT(n=18). Differentially expressed genes are statically significant, with an adjusted p-value <0,05, and are highlighted in gray.

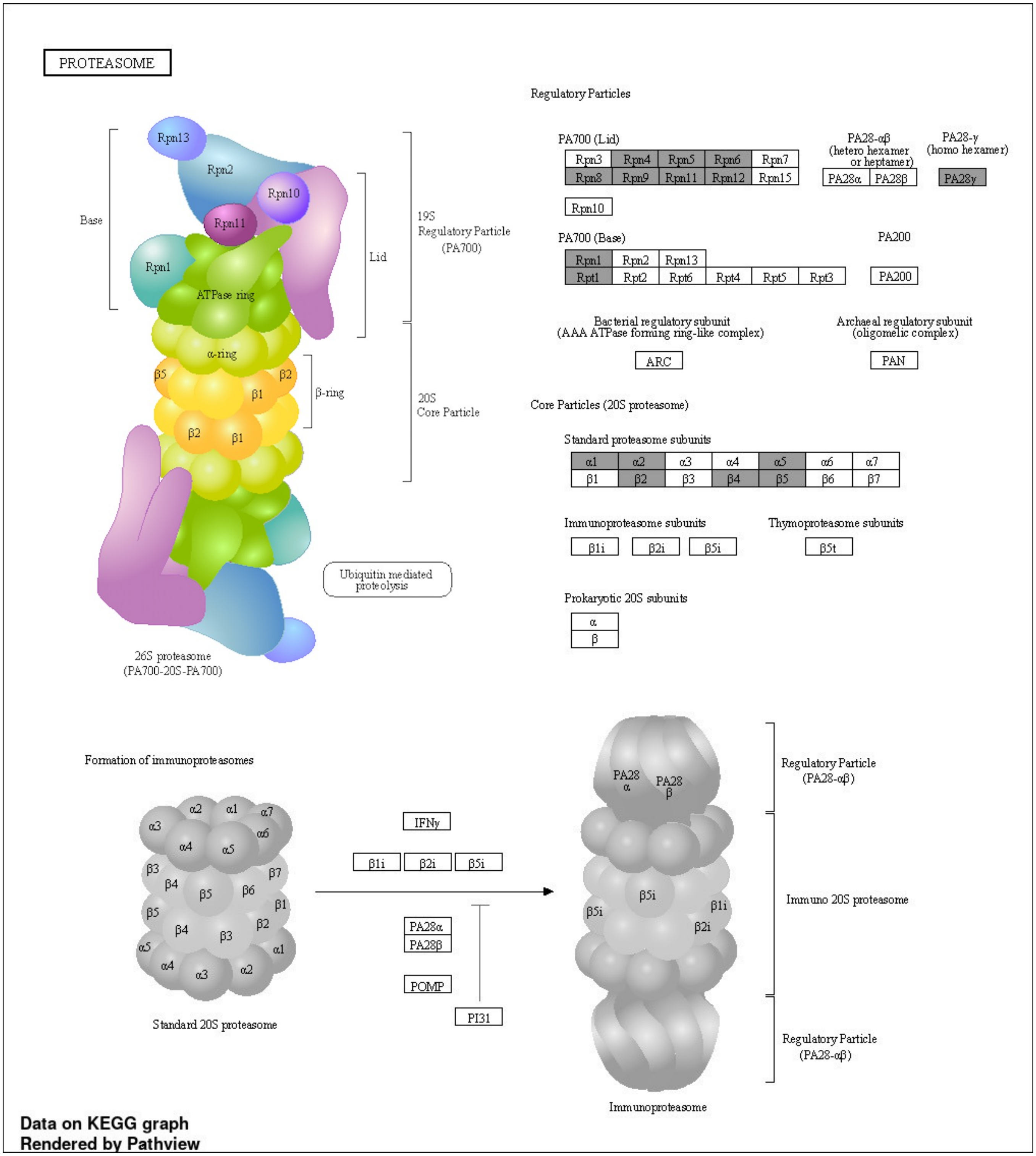


**Supplementary Figure S3. Transcriptome analysis revealed elevated mRNA expression of genes encoding proteasome subunits on stimulated naïve B cells from NFKB1 mutation carriers.** Pathway overview of elevated expression of NFKB target genes comparing NFKB1 mutation carriers (3 affected and 7 unaffected) to WT (n=18). Differentially expressed genes are statically significant, with an adjusted p-value <0.05, and are highlighted in gray.

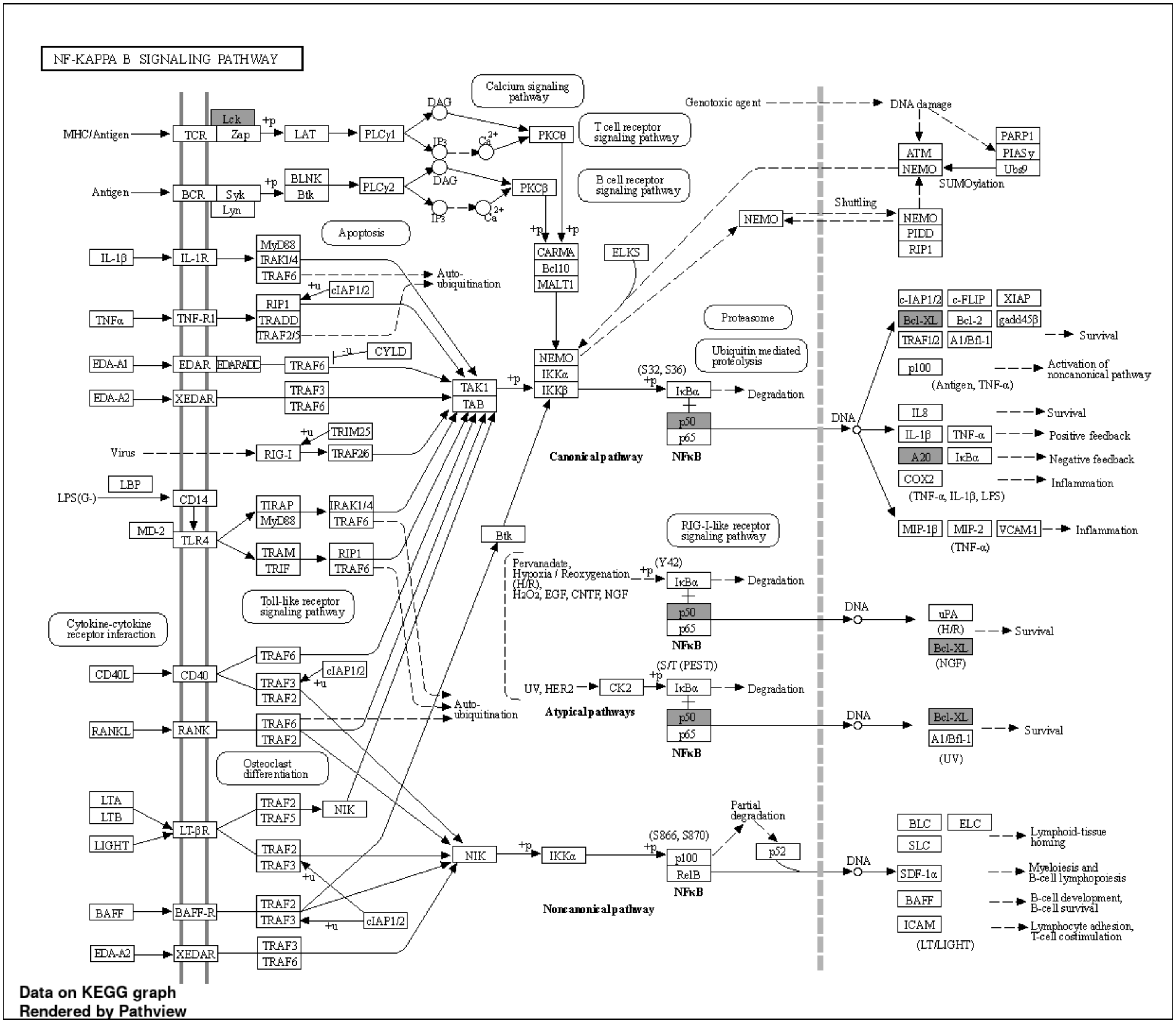


**Supplementary Figure S4**. **Transcriptome analysis revealed decreased expression of genes encoding components of the NF-kB signaling pathway on stimulated naïve B cells from unaffected NFKB1 mutation carriers.** Pathway overview of decreased expression of NFKB target genes comparing unaffected NFKB1 mutation carriers (n=7) to WT (n=18). Differentially expressed genes are statically significant, with an adjusted p-value <0.05, and are highlighted in gray.

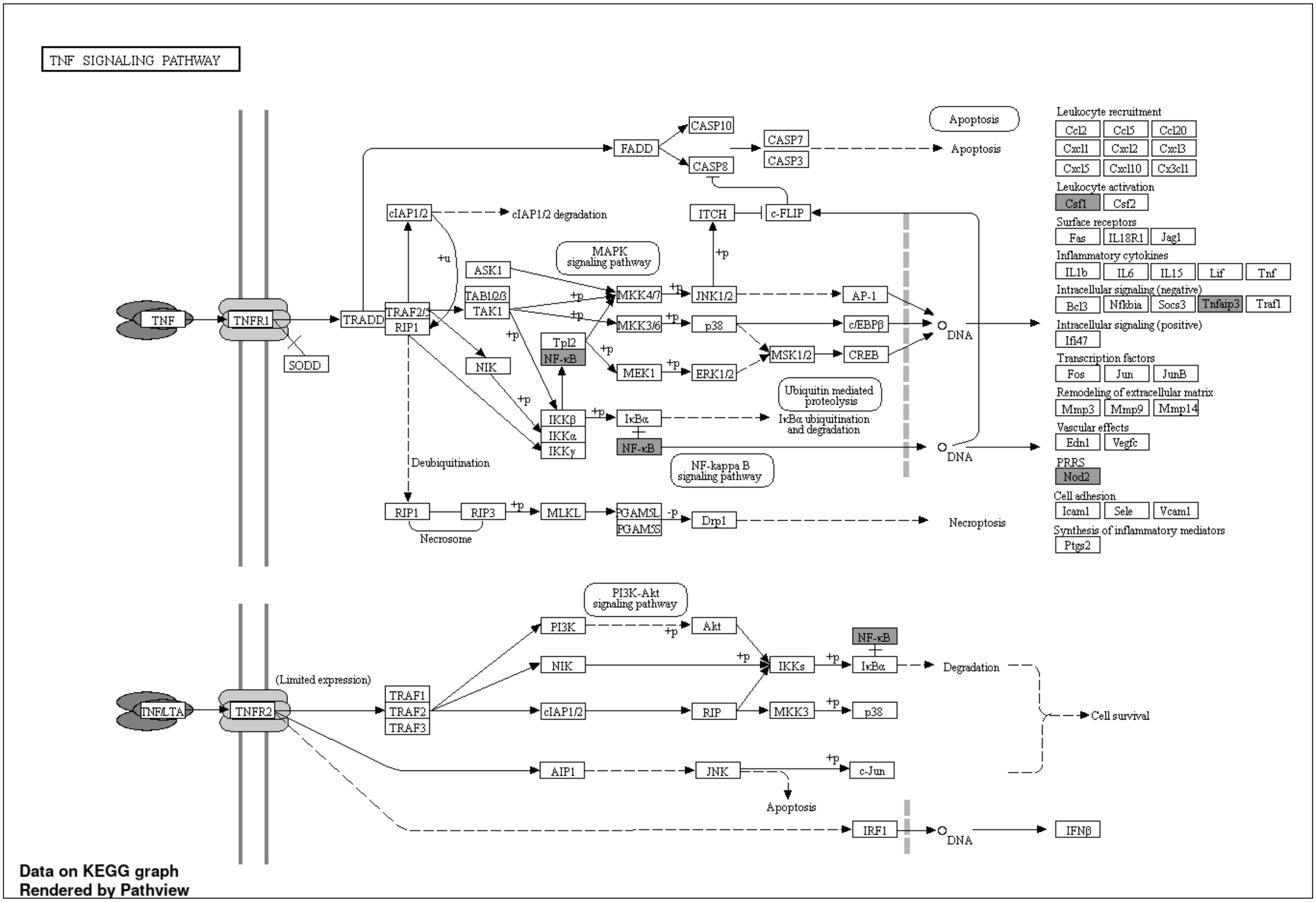


**Supplementary Figure S5**. **Reduced expression of genes encoding components of the TNF signaling pathway on stimulated naïve B cells from unaffected NFKB1 mutation carriers.** Pathway overview of NFKB target genes with decreased expression when comparing unaffected NFKB1 mutation carriers (n=7) to WT (n=18). Differentially expressed genes are statically significant, with an adjusted p-value <0.05, and are highlighted in gray.

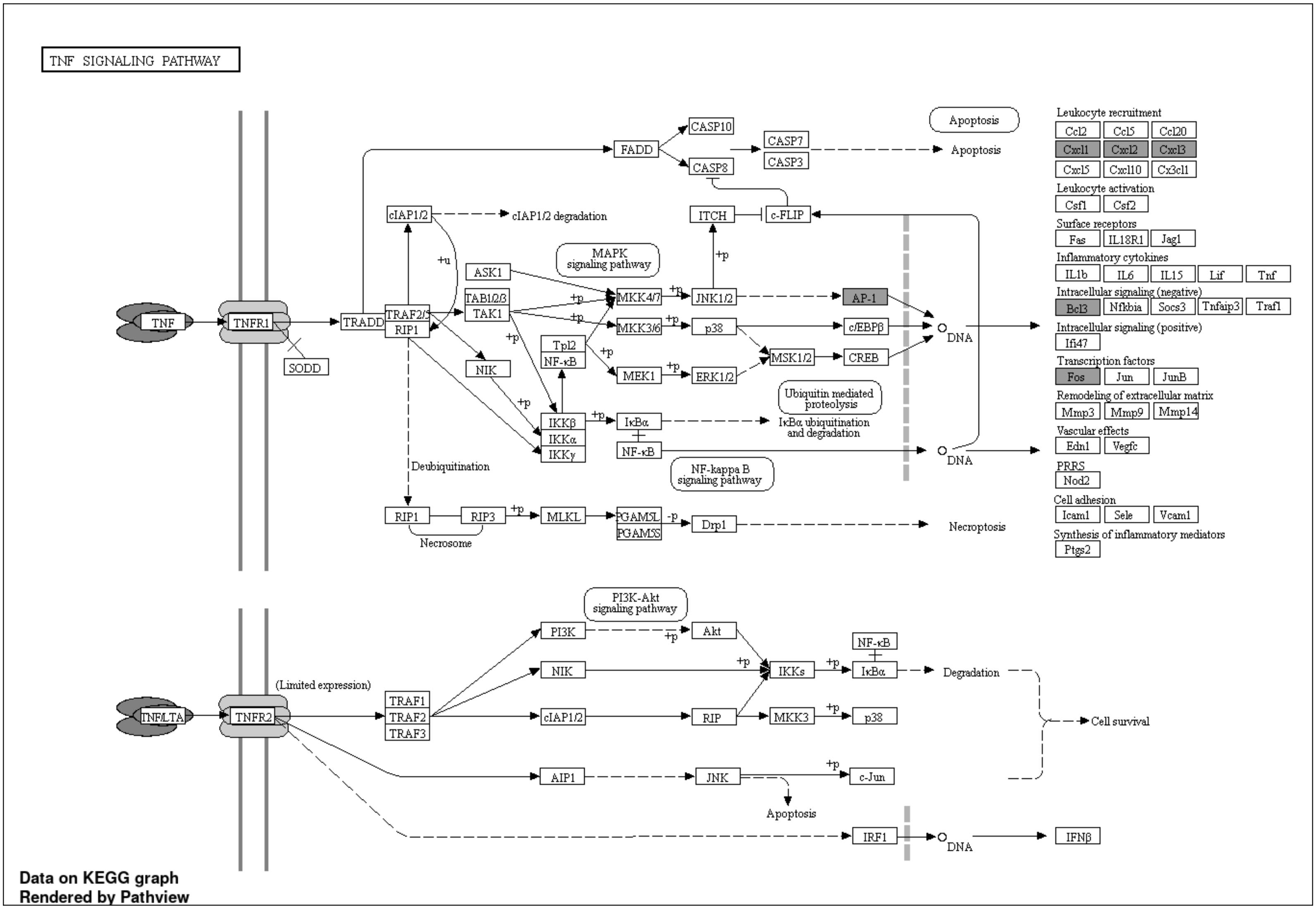


**Supplementary Figure S6**. **Increased expression of genes encoding components of the TNF signaling pathway on stimulated naïve B cells from unaffected NFKB1 mutation carriers.** Pathway overview of NFKB target genes with increased expression when comparing unaffected NFKB1 mutation carriers (n=7) to WT (n=18). Differentially expressed genes are statically significant, with an adjusted p-value <0.05, and are highlighted in gray.

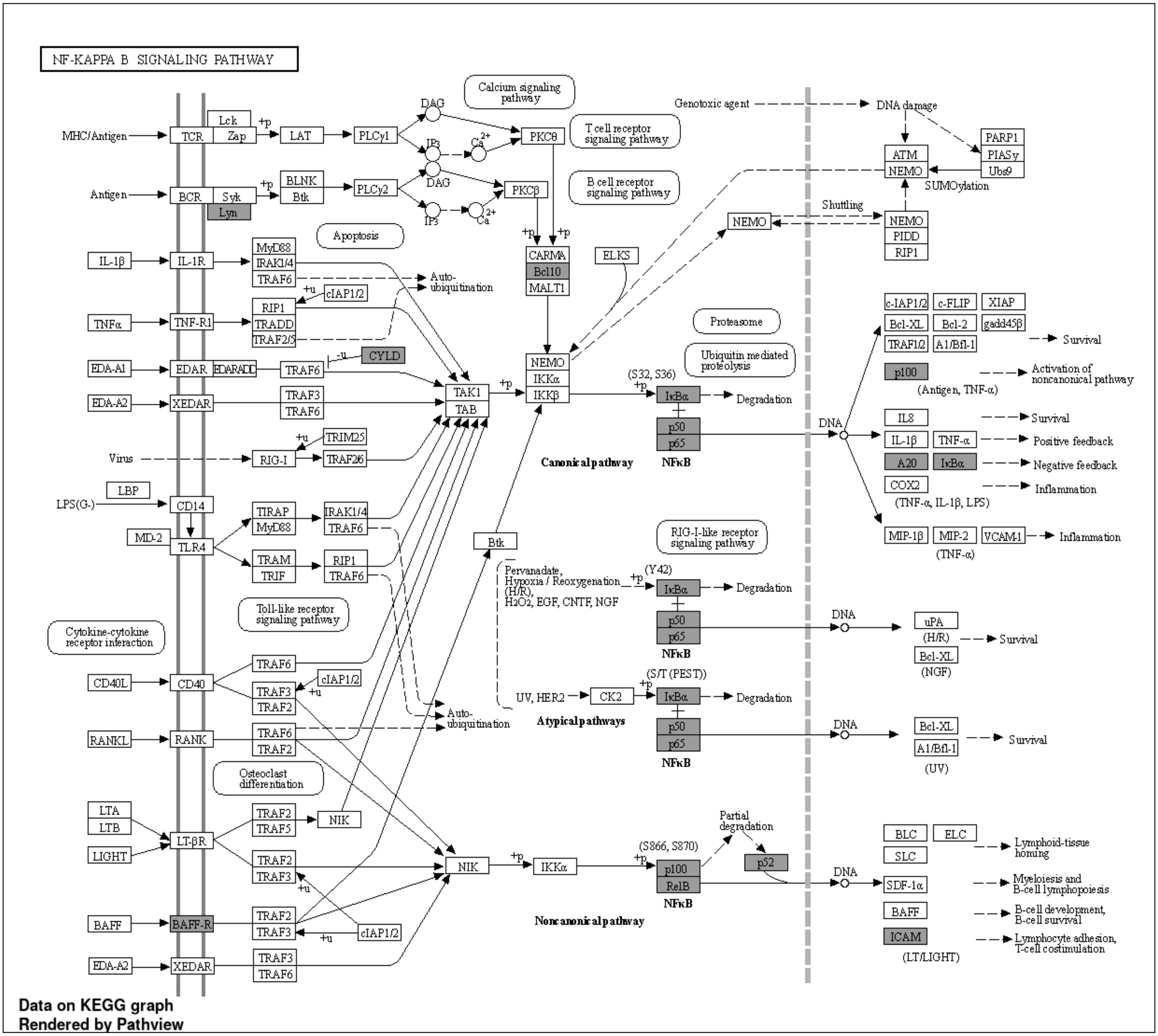


**Supplementary Figure S7**. **Transcriptome analysis revealed decreased expression of genes encoding components of the NF-kB signaling pathway on unstimulated naïve B cells from NFKB1 mutation carriers.** Pathway overview of NFKB target genes with decreased expression, when comparing unstimulated naïve B cells from NFKB1 mutation carriers (n=3) to WT (n=4). Differentially expressed genes are statically significant, with an adjusted p-value <0.05, and are highlighted in gray.

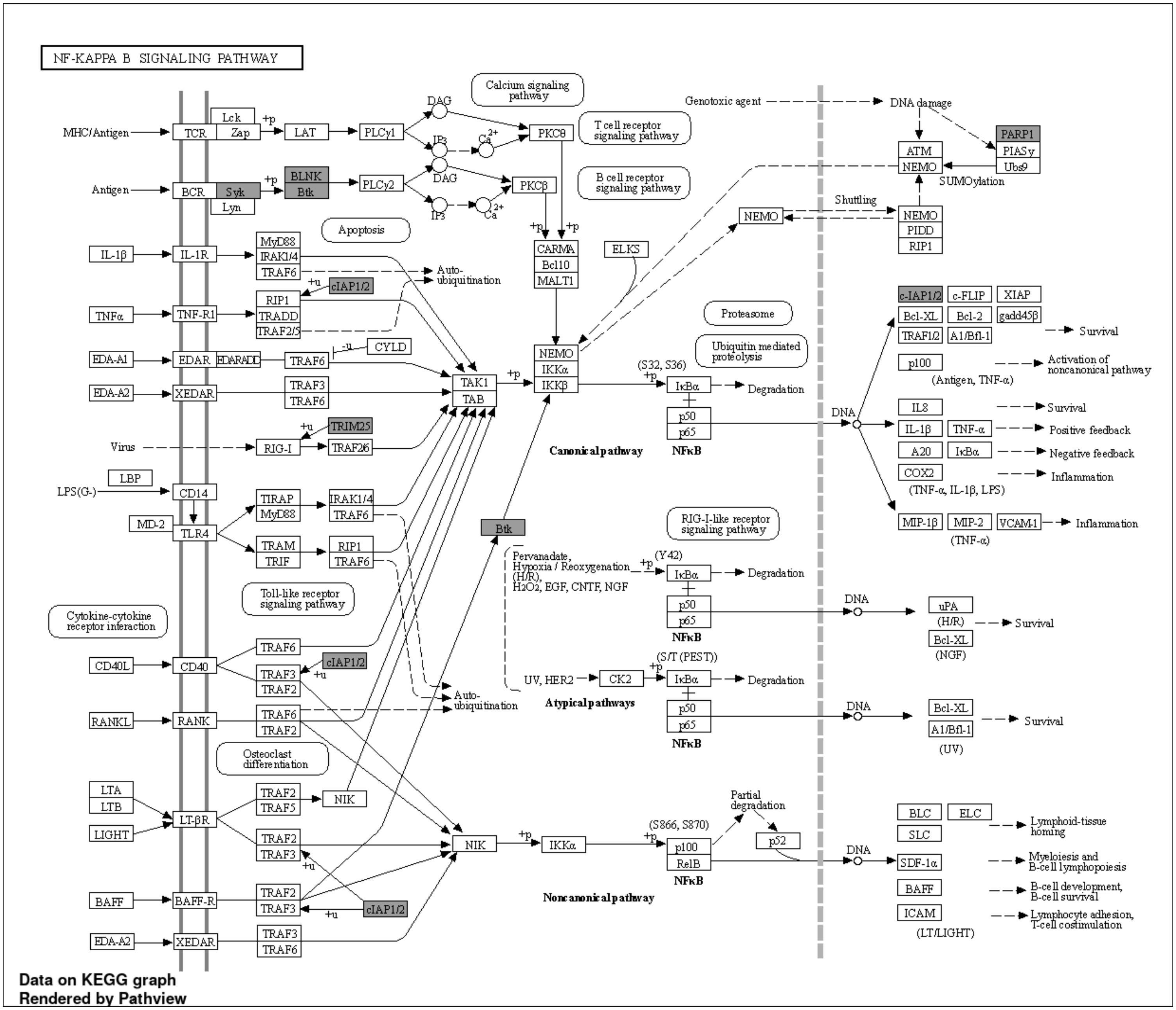


**Supplementary Figure S8**. **Transcriptome analysis revealed increased expression of genes encoding components of the NF-kB signaling pathway on unstimulated naïve B cells from NFKB1 mutation carriers.** Pathway overview of NFKB target genes with increased expression, when comparing unstimulated naïve B cells from NFKB1 mutation carriers (n=3) to WT (n=4). Differentially expressed genes are statically significant, with an adjusted p-value <0.05, and are highlighted in gray.

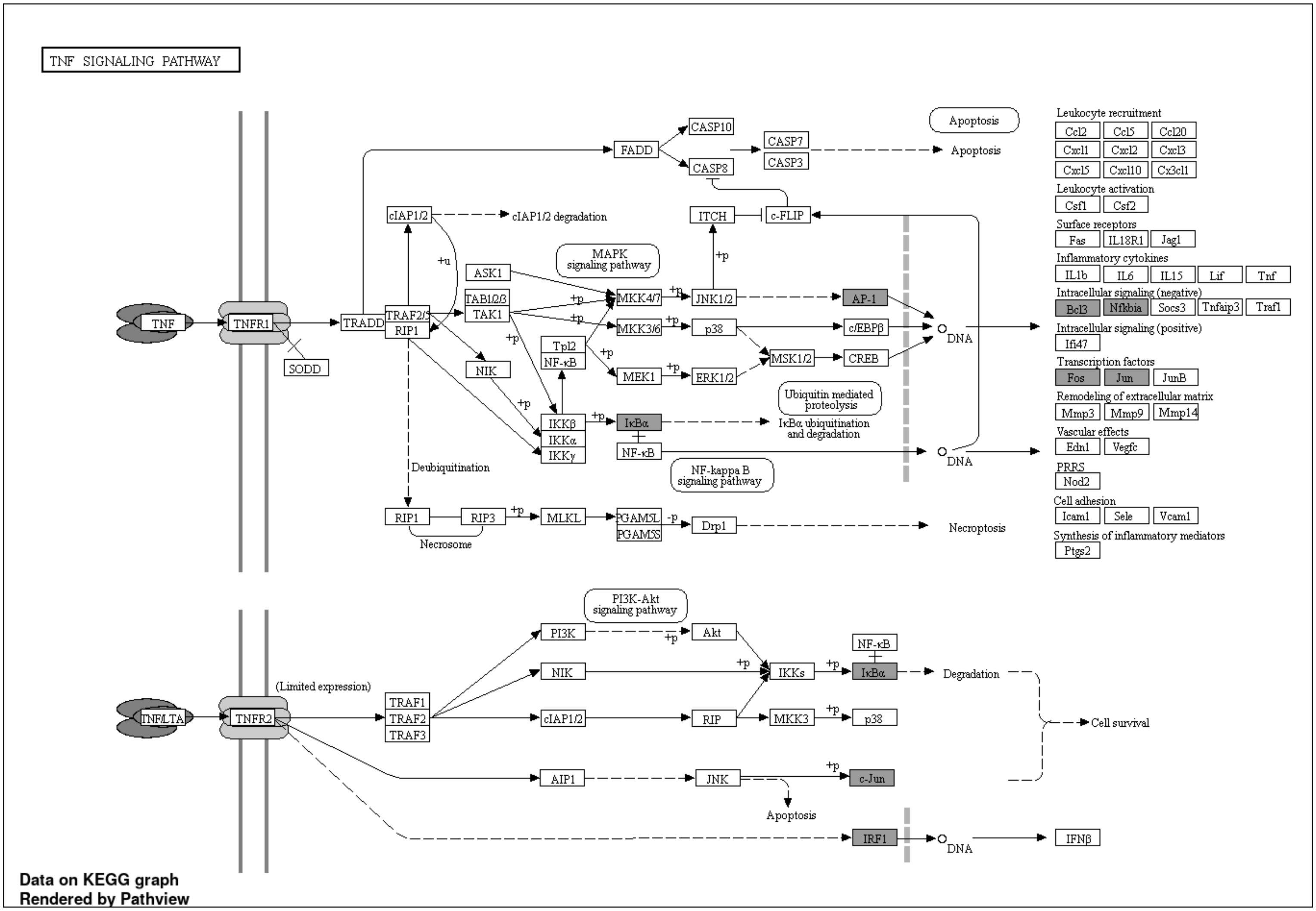


**Supplementary Figure S9**. **Transcriptome analysis revealed decreased expression of genes encoding components of the TNF signaling pathway on IgM memory B cells from NFKB1 mutation carriers.** Pathway overview of NFKB target genes with decreased expression, when comparing IgM memory B cells from NFKB1 mutation carriers (n=3) to WT (n=4). Differentially expressed genes are statically significant (p-value <0.05), and are highlighted in gray.

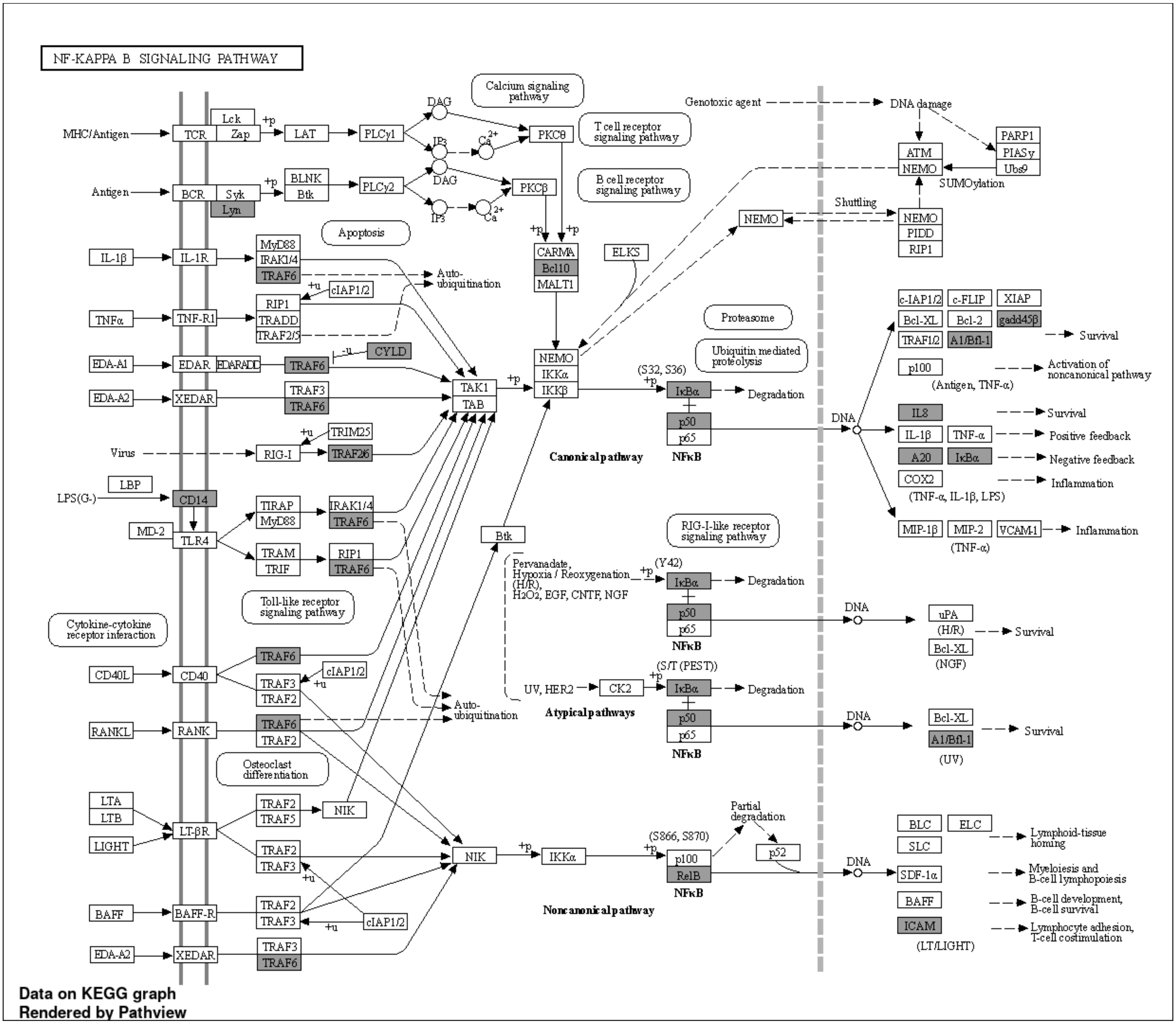


**Supplementary Figure S10. Transcriptome analysis revealed decreased expression of genes encoding components of the NF-kB signaling pathway on unstimulated switched memory B cells from NFKB1 mutation carriers.** Pathway overview of NFKB target genes with decreased expression, when comparing unstimulated switched memory B cells from NFKB1 mutation carriers (n=3) to WT (n=4). Differentially expressed genes are statically significant (p-value <0.05), and are highlighted in gray.

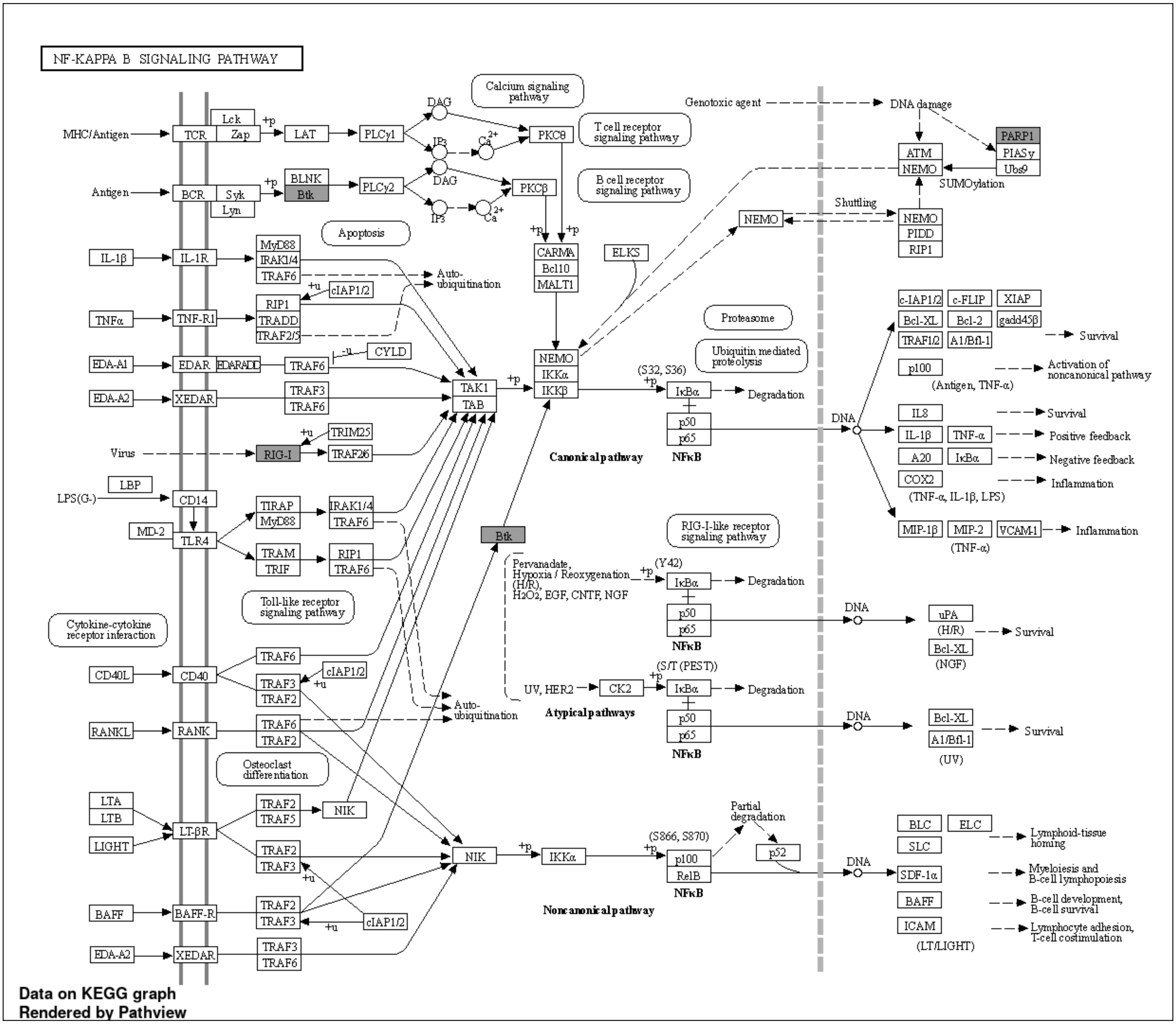


**Supplementary Figure S11**. **Transcriptome analysis revealed increased expression of genes encoding components of the NF-kB signaling pathway on unstimulated switched memory B cells from NFKB1 mutation carriers.** Pathway overview of NFKB target genes with increased expression, when comparing unstimulated switched memory B cells from NFKB1 mutation carriers (n=3) to WT (n=4). Differentially expressed genes are statically significant (p-value <0.05), and are highlighted in gray.

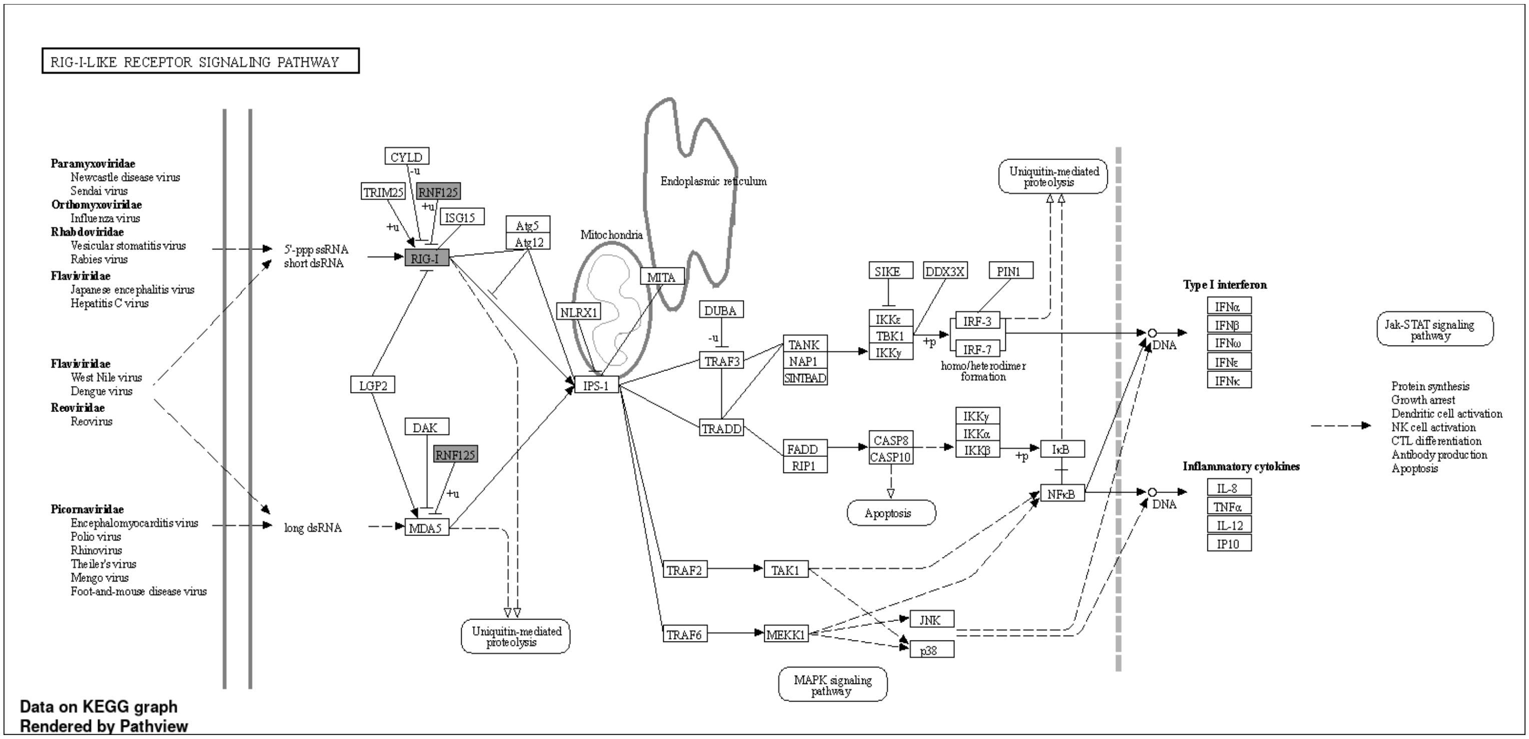


**Supplementary Figure S12**. **Transcriptome analysis revealed decreased expression of genes encoding components of the RIG-I-like receptor signaling pathway on unstimulated CD21 low B cells from NFKB1 mutation carriers.** Pathway overview of NFKB target genes with decreased expression, when comparing unstimulated CD21 low B cells from NFKB1 mutation carriers (n=3) to WT (n=4). Differentially expressed genes are statically significant (p-value <0.05), and are highlighted in gray.

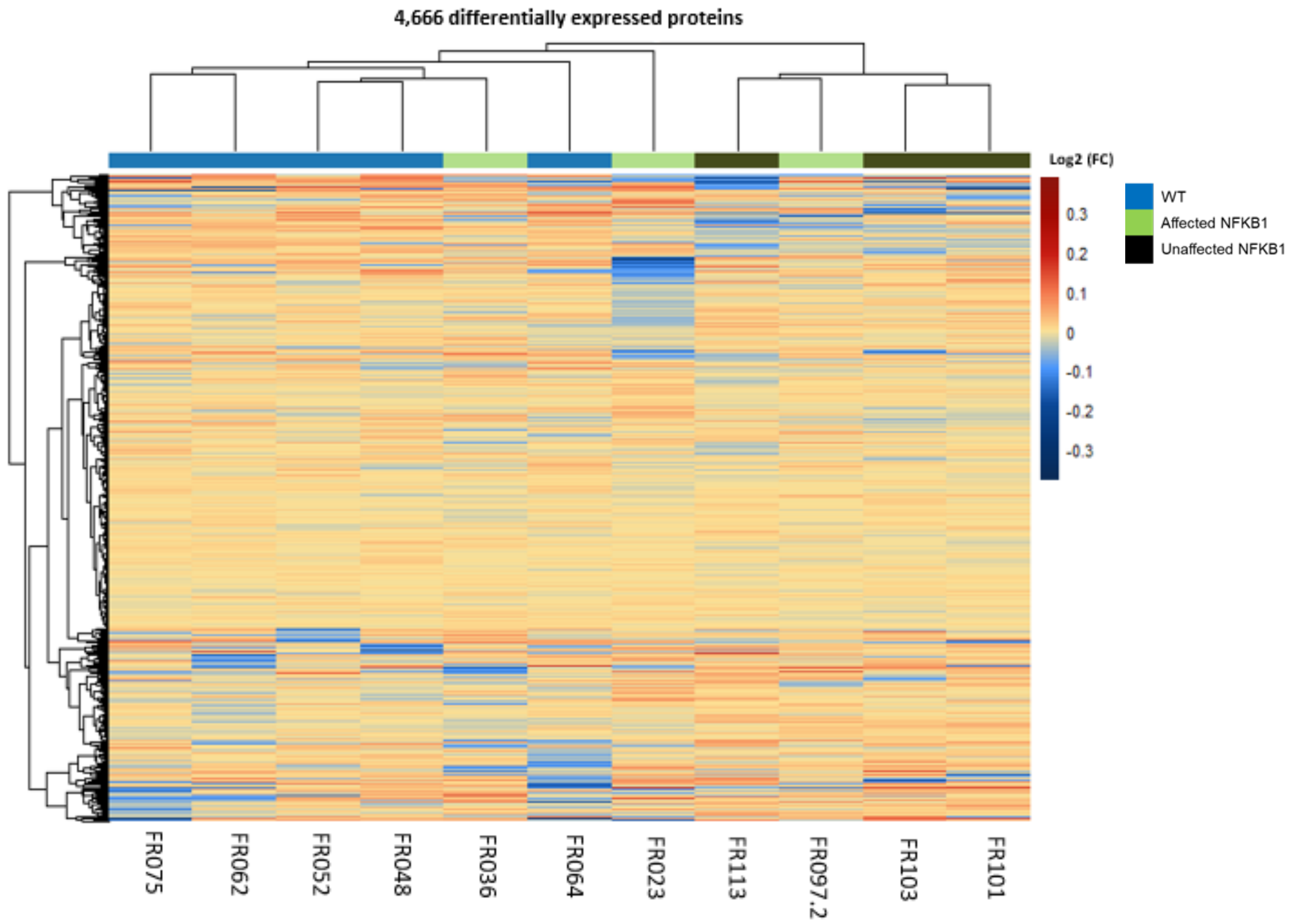


**Supplementary Figure S13. Clustering of proteome of NFKB1 mutation carriers and WT displayed as a heatmap.** Heatmap represents the hierarchical clustering of differentially expressed proteins across NFKB1 mutation carriers (n=3 affected and 3 unaffected) when compared to WT (n=5). Color indicates log fold change were blue and red represent a decrease, and increased expression, respectively. Differentially expressed proteins were identified with alpha=0.05 and absolute log2 fold change of 1.5.

Methylation and RNA-seq
n=10 NFKB1 (3 affected and 7 unaffected) vs 18 WT

ATAC-seq
n = 8 NFKB1 (3 affected-+ 5 unaffected) vs 17 WT

Proteomics
n= 7 (3 affected+ 4 unaffected) vs 11 WT

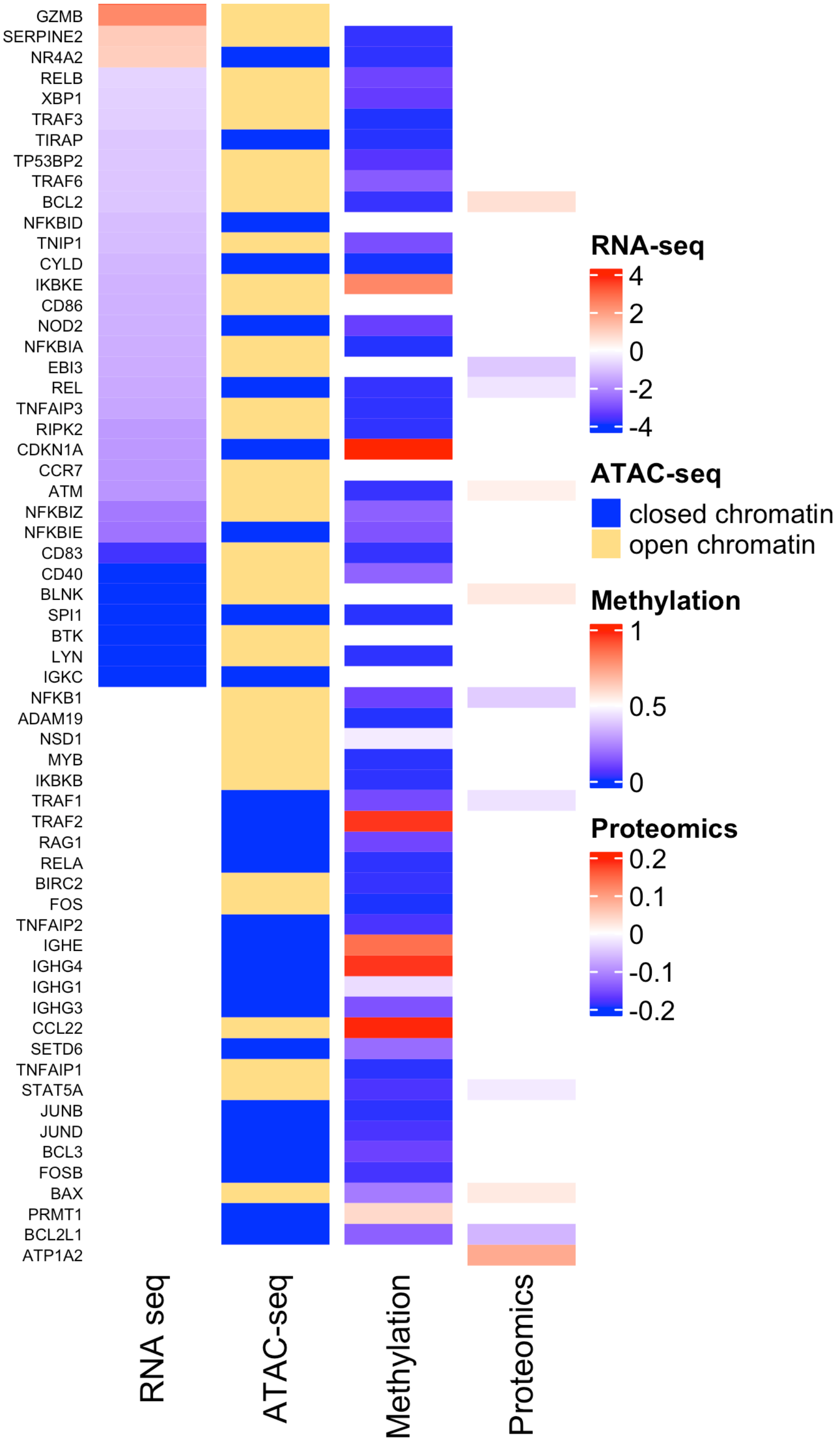


**Supplementary Figure S14.Association analysis of transcriptome, chromatin accessibility, methylation and proteomics on all NFBK1-mutated cells indicates decreased expression of NFKB target genes.** Heatmap representation showing association between transcriptome, chromatin accessibility, methylome, and proteome from NFKB1 mutation carriers compared to WT on stimulated naïve B cells. For RNA-seq data, differentially expressed genes obtained from the comparison between NFKB1 mutation carriers to WT are shown. Color indicates log fold change were blue and red represent a decrease, and increased expression, respectively. The cutoff for differential expression was set at p<0.05 adjusted p value and 0.5 for log2 fold change. For ATAC-seq, preferentially open or closed regions from *NFKB1* mutation carriers are shown. Blue bars represent preferentially closed and yellow bars preferentially open chromatin regions. Methylation data from NFKB1 mutation carriers is depicted. The median methylation is represented as a beta-value from 0 to 1 (0 to 100% methylation) at promoter regions. Color represents methylation values, where red is 100% and blue 0% methylation, respectively. For proteomic data, protein ratios were log-2 transformed. Differentially expressed proteins (alpha = 0.05 and an absolute log2 fold change of 1.5) from the comparison between NFKB1 mutation carriers and WT are represented. Color represents log fold change, where blue and red represent a decreased and increased expression, respectively

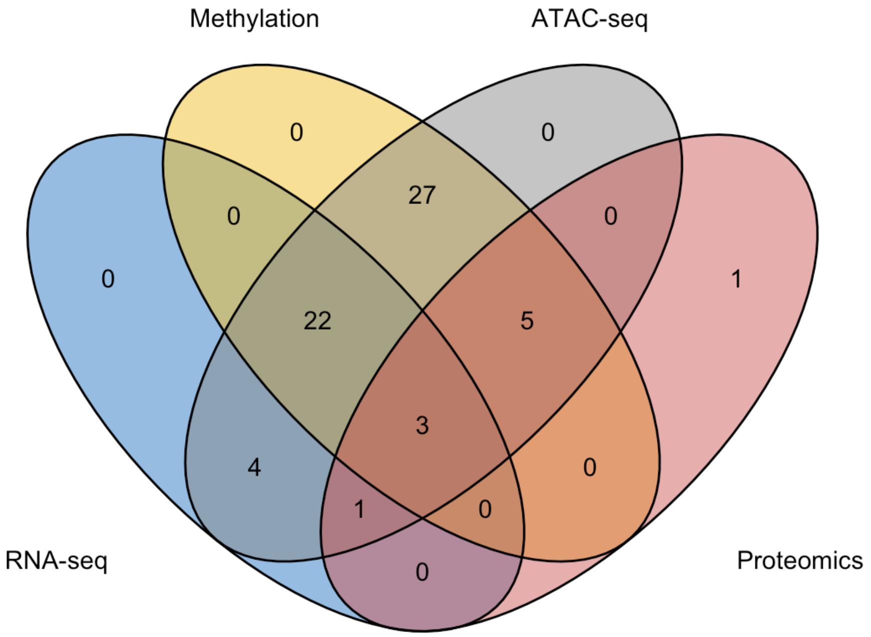


Methylation and RNA-seq
n=10 NFKB1 (3 affected and 7 unaffected) vs 18 WT

ATAC-seq
n = 8 NFKB1 (3 affected-+ 5 unaffected) vs 17 WT

Proteomics
n= 7 (3 affected+ 4 unaffected) vs 11 WT

| n=4 |
|---|
| BTK |
| CCR7 |
| GZMB |
| NFKBID |

| n=22 |
|---|
| CD83 |
| CDKN1A |
| CYLD |
| IKBKE |
| IRF2 |
| LYN |
| NFKBIA |
| NFKBIE |
| NFKBIZ |
| NOD2 |
| NR4A2 |
| RELB |
| RIPK2 |
| SERPINE2 |
| SPI1 |
| TIRAP |
| TNFAIP3 |
| TNIP1 |
| TP53BP2 |
| TRAF3 |
| TRAF6 |
| XBP1 |

| n=27 |
|---|
| ADAM19 |
| BCL3 |
| BIRC2 |
| FOS |
| FOSB |
| IGHE |
| IGHG1 |
| IGHG3 |
| IGHG4 |
| IKBKB |
| IRF1 |
| ITCH |
| JUNB |
| JUND |
| MYB |
| NSD1 |
| PRMT1 |
| RAG1 |
| RELA |
| SETD6 |
| SOCS1 |
| TAX1BP1 |
| TNFAIP1 |
| TNFAIP2 |
| TRAF2 |
| TWIST1 |
| TWIST2 |

| n=1 |
|---|
| EBI3 |

| n=3 |
|---|
| ATM |
| BCL2 |
| REL |

| n= 5 |
|---|
| BAX |
| BCL2L1 |
| NFKB1 |
| STAT5A |
| TRAF1 |

| n=1 |
|---|
| ATP1A2 |

**Supplementary Figure S15. Overlap of NFKB target genes between different data sets when comparing stimulated naïve B cells from NFKB1 mutation carriers to WT.** Venn diagram showing a supervised analysis of NFKB target genes on stimulated naïve B cells from NFKB1 mutation carriers. Diagram represents overlap between differentially expressed genes and differentially expressed proteins comparing NFKB1 mutation carriers to WT, plus preferentially open chromatin regions and methylation levels from unaffected NFKB1 mutation carriers. Associated gene names are shown in the columns below the diagram.

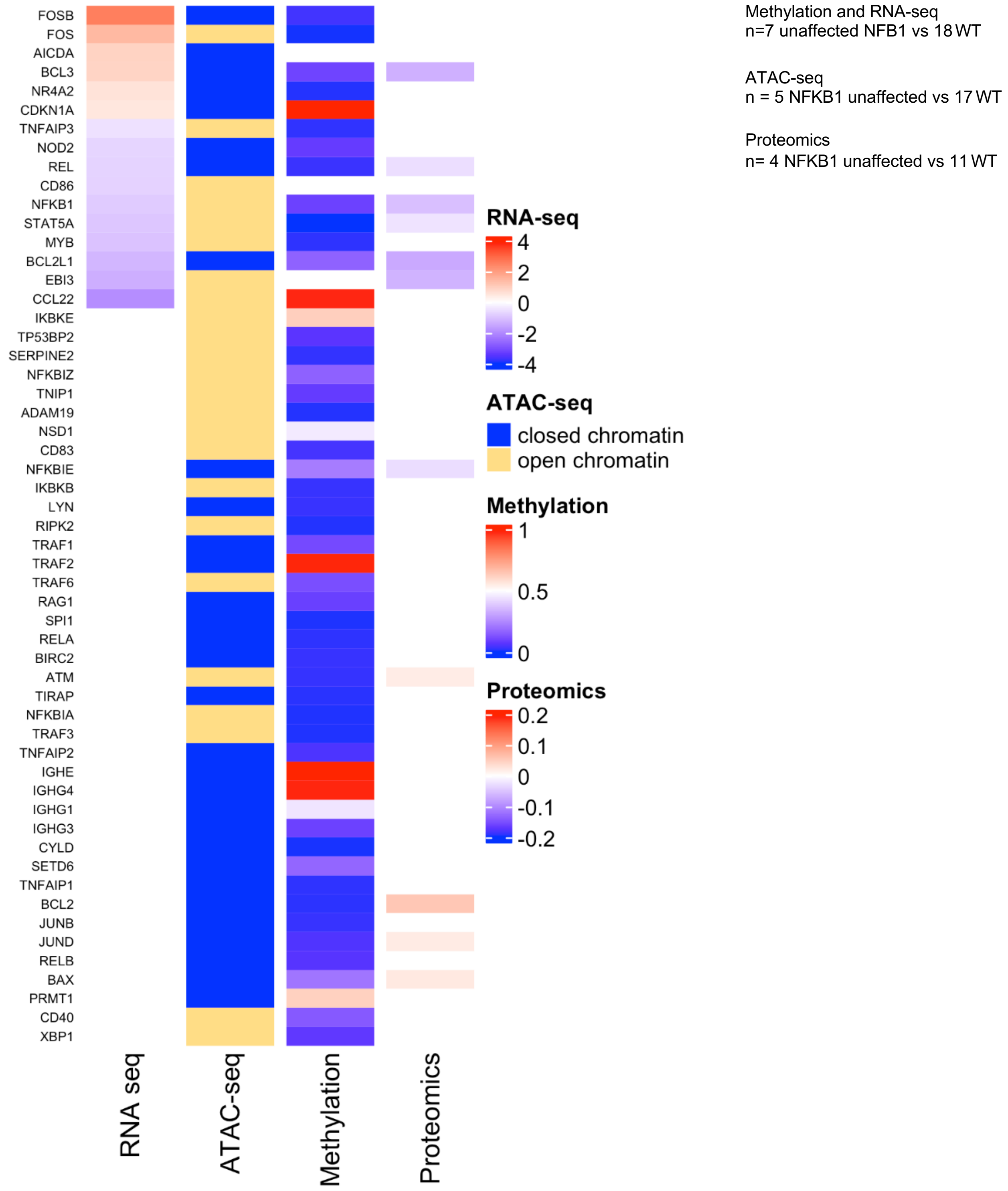


**Supplementary Figure S16. Association analysis of transcriptome, chromatin accessibility, methylation and proteomics on stimulated naïve B cells from unaffected NFKB1 mutation carriers shows predominantly reduce expression of NFKB target genes.** Heatmap representation showing association between transcriptome, chromatin accessibility, methylome, and proteome from unaffected NFKB1 mutation carriers compared to WT on stimulated naïve B cells. For RNA-seq data, differentially expressed genes obtained from the comparison between unaffected NFKB1 mutation carriers to WT are shown. Color indicates log fold change were blue and red represent a decrease, and increased expression, respectively. The cutoff for differential expression was set at p<0.05 adjusted p value and 0.5 for log2 fold change. For ATAC-seq, preferentially open or closed regions from unaffected *NFKB1* mutation carriers are shown. Blue bars represent preferentially closed and yellow bars preferentially open chromatin regions. Methylation data from unaffected NFKB1 mutation carriers is depicted. The median methylation is represented as a beta-value from 0 to 1 (0 to 100% methylation) at promoter regions. Color represents methylation values, where red is 100% and blue 0% methylation, respectively. For proteomic data, protein ratios were log-2 transformed. Differentially expressed proteins (alpha = 0.05 and an absolute log2 fold change of 1.5) from the comparison between unaffected NFKB1 mutation carriers and WT are represented. Color represents log fold change, where blue and red represent a decreased and increased expression, respectively

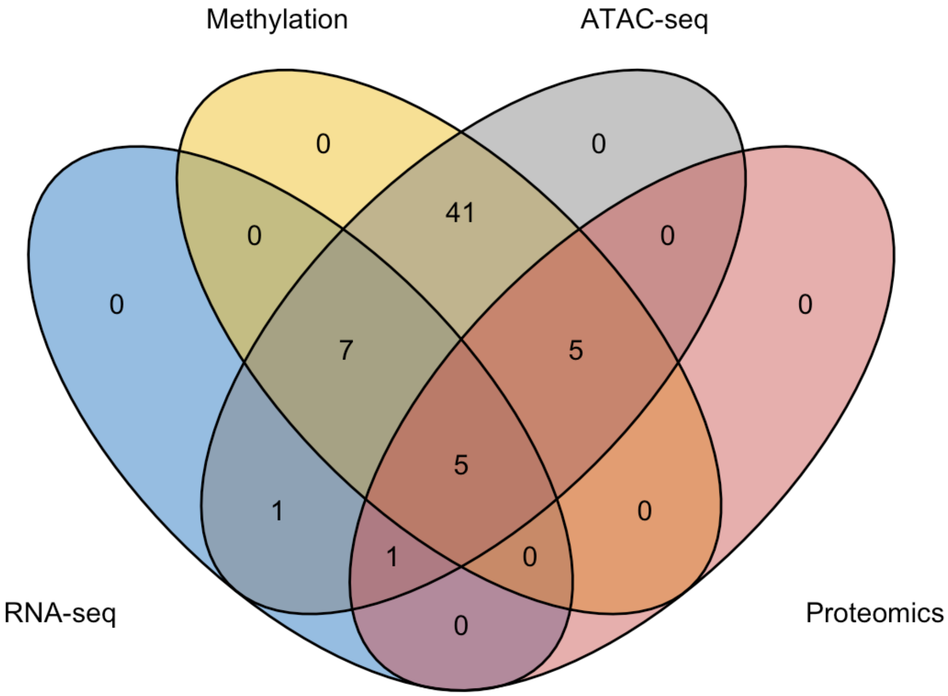


Methylation and RNA-seq
n=7 unaffected NFB1 vs 18 WT

ATAC-seq
n = 5 NFKB1 unaffected vs 17 WT

Proteomics
n= 4 NFKB1 unaffected vs 11 WT

| n=1 |
|---|
| EBI3 |

| n=7 |
|---|
| CDKN1A |
| FOS |
| FOSB |
| MYB |
| NOD2 |
| NR4A2 |
| TNFAIP3 |

| n=41 | |
|---|---|
| ADAM19 | RELB |
| BIRC2 | RIPK2 |
| CD83 | SERPINE2 |
| CYLD | SETD6 |
| IGHE | SOCS1 |
| IGHG1 | SPI1 |
| IGHG3 | TAX1BP1 |
| IGHG4 | TIRAP |
| IKBKB | TNFAIP1 |
| IKBKE | TNFAIP2 |
| IRF1 | TNIP1 |
| IRF2 | TNP2 |
| ITCH | TP53BP2 |
| JUNB | TRAF1 |
| LYN | TRAF2 |
| NFKBIA | TRAF3 |
| NFKBIZ | TRAF6 |
| NSD1 | TWIST1 |
| PRMT1 | TWIST2 |
| RAG1 | XBP1 |
| RELA | |

| n=1 |
|---|
| EBI3 |

| n=5 |
|---|
| BCL2L1 |
| BCL3 |
| NFKB1 |
| REL |
| STAT5A |

| n=5 |
|---|
| ATM |
| BAX |
| BCL2 |
| JUND |
| NFKBIE |

**Supplementary Figure S17. Overlap of NFKB target genes between different data sets when comparing stimulated naïve B cells from unaffected NFKB1 mutation carriers to WT.** Venn diagram showing a supervised analysis of NFKB target genes on stimulated naïve B cells from unaffected NFKB mutation carriers. Diagram represents overlap between differentially expressed genes and differentially expressed proteins comparing unaffected mutation carriers to WT, plus preferentially open chromatin regions and methylation levels from unaffected NFKB1 mutation carriers. Associated gene names are shown in the columns below the diagram.
